# Key metabolic pathways involved in seed primary physiological dormancy maintenance of Korean pine seed

**DOI:** 10.1101/2021.02.08.430286

**Authors:** Yuan Song, Xiaoye Gao

**Affiliations:** College of Eco-Environmental Engineering, Guizhou Minzu University, Guiyang, China 550003

## Abstract

The metabolic changes that occurred during either cold stratification or after-ripen treatment, and in both dormant seeds and after-ripened seeds either under the dry state or during imbibition have been extensively explored. Much less is known about those present in both dormant seeds and cold stratified seeds during the same period of incubation under favorable germination conditions. Metabolite composition was investigated in both embryo and megagametophyte of primary physiological dormant seeds (PPDS) of *Pinus Koreansis* collected at 0 week, 1 week, 2 weeks, 4 weeks and 6 weeks of incubation, and of cold stratified seeds with released primary physiological dormancy (RPPDS) sampled at 0 week and 1 week of incubation, seed coat rupture stage and radicle protrusion stage. Embryo contained higher levels of most metabolites compared to megagametophyte. Strong metabolic changes occurred at 1 week and 4 weeks of incubation in PPDS, with most metabolites were significantly accumulated in 4-weeks-incubated PPDS. A larger metabolic switch was found in RPPDS between 1-week-incubation and seed coat rupture stage. Especially, there was a significant major decrease in the relative levels of most phosphorylated sugars and amino acids. The carbohydrate metabolism, especially pentose phosphate pathway and tricarboxylic acid cycle were more active pathways in the embryos of 4-weeks-incubated PPDS, but the operation rate of most amino acid metabolism was lower compared to 1-week-incubated RPPDS. We suggest that a larger metabolic switch in the embryo of PPDS after 4 weeks of incubation may assist in maintaining primary dormancy.

**One-sentence summary:** A larger metabolic switch in dormant seeds after 4 weeks of incubation under favorable conditions for germination may maintain primary physiological dormancy of Korean pine seeds.

---

Woody plant species determines the structure of forest ecosystem and thus has a profound impact on the ecological conditions found within forest (Zaynab et al., 2018). Seed germination is a key stage in the plant life cycle. Non-dormant seeds germinate under favorable conditions for seedling establishment, beginning the second round of plant life cycle (He et al., 2011). Those orthodox seeds are dry after maturity (De Gara et al., 2000). Seed germination process can be divided three phases based on the characteristic of triphasic mode of water uptake (Bewley, 1997, Finch-Savage and Leubner-Metzger, 2006). A rapid and passive water uptake by dry seeds first occurs during phase I. Then, there is a slow water uptake in phase II during which metabolic activities resume, preparing for subsequent seed germination (He et al., 2011b). The occurrence of further water uptake during phase III is accompanied by the radicle protrusion through seed coat. Seed germination sensu stricto refers to the physiological process spanning from the uptake of water by dry seed to radicle protrusion (Bewley, 1997). For some species, seeds are non-dormant after dispersal from mother plant and rapidly germinate when encounter favorable conditions. However, to survive in unfavorable climates, plants have evolved a seed dormancy mechanism that allow seeds to persist for prolonged periods until conditions for seedling establishment become favorable (De et al., 2019). Seed dormancy can be released either by cold stratification (a low-temperature storage of imbibed seeds) or by after-ripening (a period of dry storage) (Bewley et al., 2012) depending on species and environmental conditions experienced by seed during maturation.

Seed germination is a very complex physiological and biochemical process that is mainly determined by genetic factors with a substantial environmental influence (Joosen et al., 2013). Previous studies have revealed that a series of physiological and biochemical process such as repair mechanisms, transcription, translation, reactivation of metabolism, reserves mobilization, redox homeostasis regulation and organellar reassembly is initiated in imbibed seeds (Bewley, 2013; He et al., 2011b). Germination is driven by these metabolic processes. The combination of high-throughput and large scale-omics methods is used to unveil the regulation mechanism of seed germination. Due to the great potential in agriculture and the availability of genome sequence data, the genomic, transcriptomic, and proteomic studies of seed germination have largely focused on model species, such as Arabidopsis (Nakabayashi et al., 2005; Holdsworth et al., 2008a, b; Narsai et al., 2017a), rice (Narsai et al., 2017b), maize (Sekhon et al., 2013), barley (Lin et al., 2014), and wheat (Yu et al., 2014). These studies revealed that one fourth of stored mRNA are metabolism-related in dry seeds, and metabolism-related genes are also highly expressed during germination (Nakabayashi et al., 2005). Metabolism-related proteins are the major groups detected in germinated seeds (He et al., 2011b; Sano et al., 2012). The central carbon metabolism pathways provide energy and building blocks for various metabolic activities (Rosental et al., 2014). It has been suggested that glycolysis, fermentation, the tricarboxylic acid (TCA) cycle and the oxidative pentose phosphate pathway (PPP) are activated during seed germination, with energy is mainly provided by anaerobic respiration, such as fermentation at the early stage of germination (He et al., 2011a, b; Yang et al., 2007). Despite recent great advances in omics studies of seed dormancy and germination, few experiments focus on metabolome-level changes during seed dormancy maintenance and germination in woody plants.

Several studies have been conducted to explore the changes in the proteome or metabolome during either cold stratification or after-ripen for the determination of metabolic activities involved in seed germination mechanisms (Lee et al., 2006; Angelovici et al., 2011; Arc et al., 2012; Pawłowski and Staszak, 2016; Chang et al., 2018). It is the metabolic activities occurred before radical protrusion that are responsible for the delay or no germination of seeds (Bewley et al., 2013). In addition, it has been previously shown that water imbibition itself can trigger some various in gene expression or the entire metabolism whatever the depth of seed dormancy (Preston et al., 2009; Yazdanpanah et al., 2017; Xia et al., 2018b). Thus, in order to determine the real physiological and biochemical process involved in seed germination regulation, it is necessary to make dormant and non-dormant seeds to imbibe water for same time during a time window in which radicle has not yet to break through the seed coat. There are also comparative proteomic, transcriptomic, or metabolomic studies generally that are performed on dormant seeds and after-ripened (AR) seeds either under the dry state or during imbibition (Cadman et al., 2006; Chibani et al., 2006; Gao et al., 2012; Joosen et al., 2013; Lu et al., 2016; Yazdanpanah et al., 2017; Xia et al., 2018b; Xu et al., 2020). The mechanisms controlling seed germination are well understood by comparing imbibed dormant and AR seeds. However, seed dormancy release of many species depends on cold stratification. While there has been no study that investigates the metabolic changes in both dormant seeds and cold stratified seeds following transfer to germinative conditions. Fait et al. (2006) have shown that the relative levels of TCA-cycle intermediates, 2-oxoglutarate and isocitrate, 3-phosphoglycerate, fructose 6 phosphate, and glucose 6 phosphate, increased during vernalization and were further enhanced during germination *sensu stricto*, but how the metabolites in dormant seeds change over imbibition time are still unknown.

It has been reported that different tissues of the seed such as embryo and endosperm, have specific functions during seed germination (Baskin and Baskin, 2004; Lee et al., 2006; Xu et al., 2016). Many of previous studies do not investigate the metabolic changes occurring in different seed tissues due to a limitation of sample separation techniques (Ferrari et al., 2009). The real differential metabolic activities between dormant and non-dormant seeds are possibly ignored given the use of mixed tissues over the period of experiment. This requires in future work distinguishing properly the roles of embryo from the endosperm in either dormancy release and subsequent germination or dormancy maintenance.

Korean pine is a dominant species in Mixed-broadleaved Korean pine forests (MKPF) in northeast China. Korean pine seeds are physiological dormancy caused by an endogenous block in the embryo. In our previous metabolome study of dormant and cold stratified Korean pine seeds, most metabolites in embryo substantially increased in both dormant and cold stratified seeds during the first 5 days of incubation under favorable conditions. After 5 days of incubation, cold stratified seeds exhibited a lager decrease in the relative levels of almost all amino acids and the TCA-cycle and glycolysis intermediates, while most of metabolites in dormant seeds were maintained at a relative steady level. In the study mentioned above, whole experiment was only carried out for 11 days making it impossible to determine if the embryo of dormant seeds continue to maintain a relatively slow and stable metabolic state with increasing incubation duration. Furthermore, it is still unclear whether those metabolites with decreased relative levels in the embryos of cold stratified seeds show a continuous decrease or a surprising increase during the period from 11 days of incubation to testa rupture and further radicle protrusion. Based on previous those results, it has been previously hypothesized that oxygen consumption would largely decrease through reducing TCA and glycolysis in the embryos of cold stratified seeds rates when compared to dormant seeds. Thus, more oxygen might be supplied to the megagametophytesand used to produce sugars, preparing for rapid germination. However, in the studies mentioned-above, the metabolic changes that occur in megagametophyte were not determined. Thus, the objective of this study was to compare metabolomic profiles of embryo and megagametophyte from dormant dry seeds and seeds with released primary dormancy during incubation under favorable conditions, and reveal key metabolic processes involved in seed dormancy maintenance and germination.

## RESULTS

### PCA of the relative contents of metabolites in PPDS

PCA was performed on PPDS or RPPDS to investigate metabolic processes at global level. The first component of PCA clearly separated embryo samples (PPDS-0-E, PPDS-1-E, PPDS-2-E, PPDS-4-E and PPDS-6-E) and megagametophyte samples (PPDS-0-M, PPDS-1-M, PPDS-2-M, PPDS-4-M and PPDS-6-M), accounting for 41.6% of the variance. PPDS-0-E and PPDS-0-M samples formed two more discrete groups on the first principal component (Fig. 1A). The transition from 1 WAI (week after incubation) to 2 WAI stage was associated with smaller differences in the relative of contents of metabolites in both embryos and megagametophytes. The samples collected at 4 WAI and 6 WAI were also situated relatively closely to one another. The metabolome of both embryo and megagametophyte in PPDS started to deviate at 4 WAI on the second principal component.

**Figure 1.**
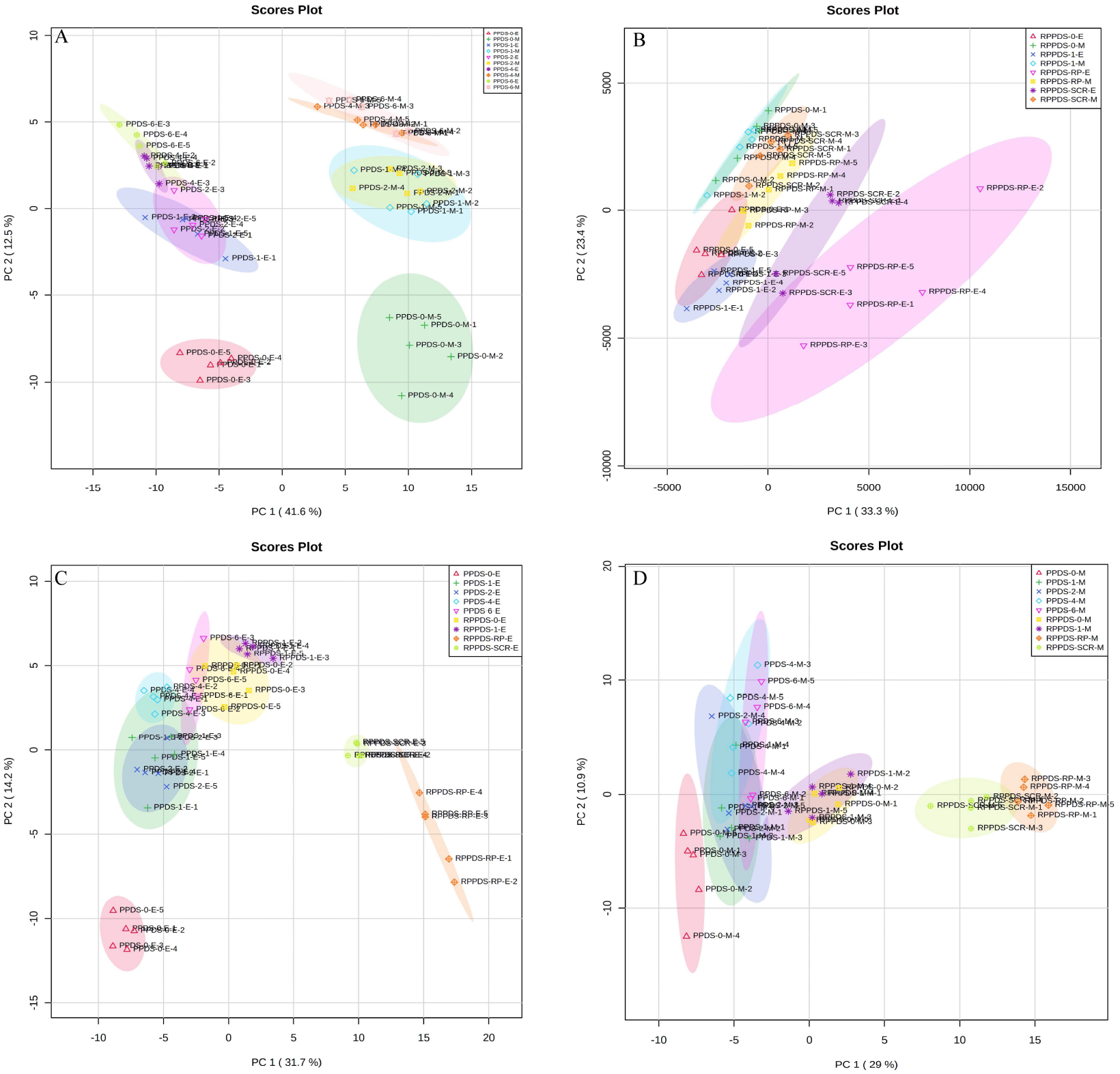
Principal components analysis (PCA) of metabolite profiles of (A) the embryos of five primary physiological dormancy seeds (PPDS) samples (PPDS-0-E, PPDS-1-E, PPDS-2-E, PPDS-4-E and PPDS-6-E) and the megagametophytes of five PPDS samples (PPDS-0-M, PPDS-1-M, PPDS-2-M, PPDS-4-M and PPDS-6-M), (B) the embryos of four seeds with released primary physiological dormancy (RPPDS) samples (RPPDS-0-E, RPPDS-1-E, RPPDS-SCR-E and RPPDS-RP-E) and the megagametophytes of four RPPDS samples (RPPDS-0-M, RPPDS-1-M, RPPDS-SCR-M and RPPDS-RP-M), (C) the embryos of PPDS (PPDS-0-E, PPDS-1-E, PPDS-2-E, PPDS-4-E and PPDS-6-E) and RPPDS (RPPDS-0-E, RPPDS-1-E, RPPDS-SCR-E and RPPDS-RP-E), and (D) the megagametophytes of PPDS (PPDS-0-M, PPDS-1-M, PPDS-2-M, PPDS-4-M and PPDS-6-M) and RPPDS (RPPDS-0-M, RPPDS-1-M, RPPDS-SCR-M and RPPDS-RP-M).

### PCA of relative contents of metabolites in RPPDS

RPPDS embryo experienced relatively larger metabolome changes than megagametophyte during incubation (Fig. 1B). Embryo did not display distinct alterations in metabolome after 1 week of incubation. However, there was a large metabolic shift after seed coat rupture, indicatedby the apparent separation between RPPDS-1-E and RPPDS-SCR-E samples along the first principal component. A relatively small difference was found between RPPDS-1-M and RPPDS-SCR-E samples.

### PCA of the relative contents of metabolites in the embryo and megagametophyte of both PPDS and RPPDS

PPDS-4-E sample was clearly differentiated from RPPDS-1-E, RPPDS-SCR-E and RPPDS-RP-E samples on the first dimension of the PCA (Fig. 1C). A similar distinction was also found between RPPDS-1-M, RPPDS-SCR-M, RPPDS-RP-M and PPDS-4-M samples (Fig. 1D).

### Fold changes of important metabolites with a VIP value > 1 and *P* < 0.05 during incubation of PPDS

When comparing the patterns of change in the relative contents of metabolites in embryos with VIP >1 and *P* < 0.05 between successive time points, a large variation occurred in the first week (43 increased, 44 decreased) (Fig. 2A), with few changes observed between 1 and 2 WAI (4 increased, 16 decreased) (Fig. 2B) and between 4 and 6 WAI (20 increased, 14 decreased) (Fig. 2D). There was a relatively higher alternation between 2 and 4 WAI (45 increased, 14 decreased) (Fig. 2C). Those metabolites that displayed a major increase in their relative contents after 1 week of incubation were mainly involved in major carbohydrate metabolism. 6-phospho-D-gluconate, D-mannose 1-phosphate, 3-phospho-D-glycerate, Beta-D-fructose-6-phosphate, phosphoenolpyruvate, ribulose 5-phosphate, glycerol 3-phosphate, D-fructose 1,6-bisphosphate, fructose 1-phosphate increased 5-to 23-fold between 0 and 1 WAI (Fig. 2A). Of PPP intermediates, D-ribose 5-phosphate also increased after 2 WAI (Fig. 2B). In contrast, several sugar alcohols, including ribitol, D-sorbitol, myo-inositol and maltitol exhibited a significant decline in their relative levels at the 1 WAI time point (Fig. 2A). The relative amounts of uridine 5’-diphosphate, cytidine 5’-monophosphate, uridine diphosphate glucose, s-methyl-5’-thioadenosine and uridine 5’-monophosphate were significant lower in 2-weeks-incubated PPDS as compared to 4-weeks-incubated PPDS (Fig. 2C). The abscisic acid relative content did not change significantly during the first 2 weeks of incubation, and strongly increased 2-fold after 4 weeks (Fig. 2C). Two TCA-cycle intermediates (citrate and cis-aconitate) and three phosphorylated sugars including D-fructose 1,6-bisphosphate, D-ribulose 5-phosphate and alpha-D-galactose 1-phosphate were also significantly accumulated from 2 WAI to 4 WAI (Fig. 2C). Moreover, the relative level of citrate, both the number and relative contents of phosphorylated sugars also increased in 4-weeks-incubated PPDS when compared to 1-weeks-incubated PPDS (Fig. 2E). Only 15 metabolites with VIP > 1 and *P* < 0.05 exhibited large differences between 1 WAI and 6 WAI (Fig. 2F).

**Figure 2.**
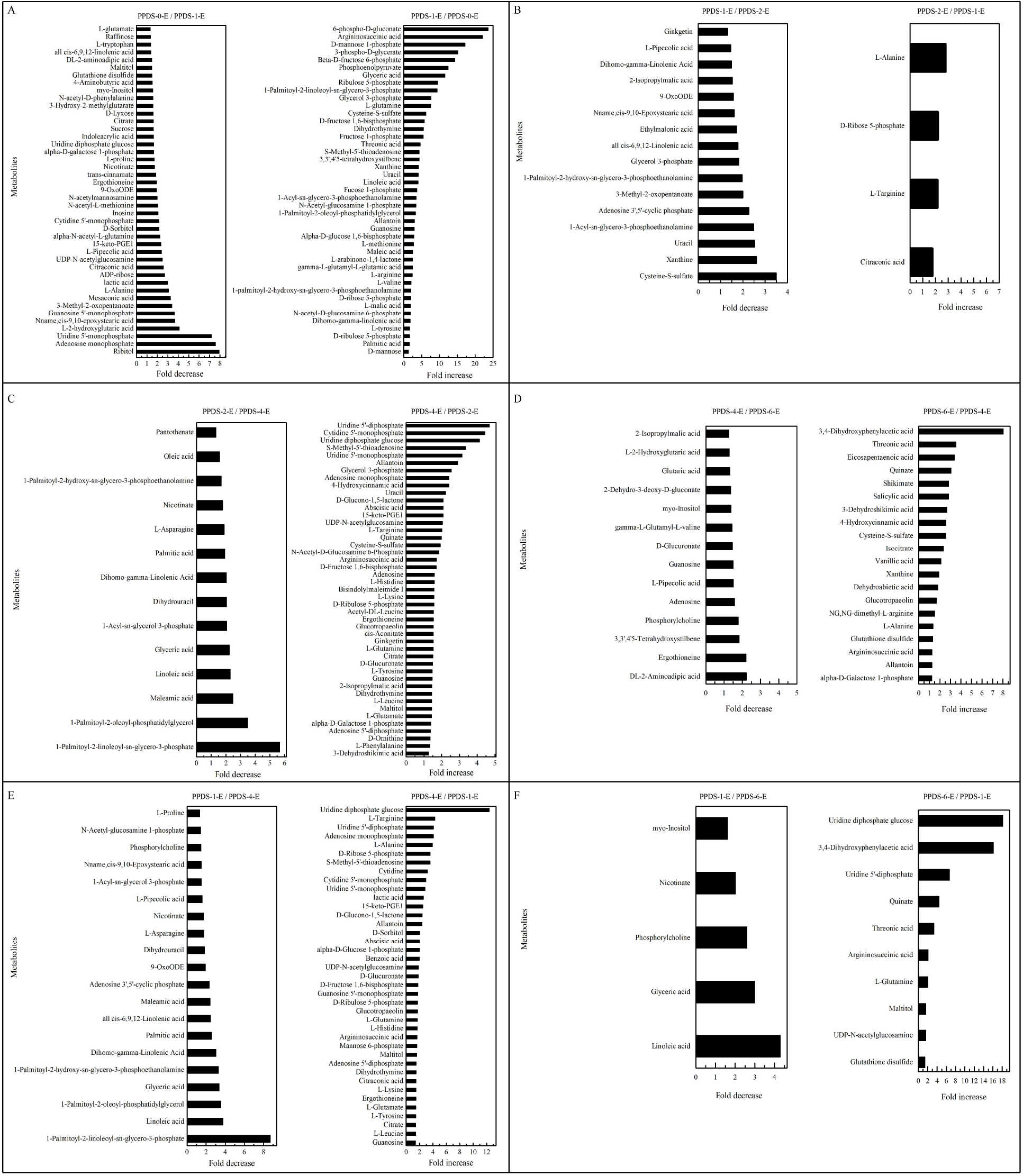
Fold change in the relative contents of important metabolites with VIP value > 1 and *P* < 0.05 in the embryos of primary physiological dormancy seeds (PPDS) (A) between 0 and 1 week of incubation (PPDS-0-E vs. PPDS-1-E), (B) 1 and 2 weeks of incubation (PPDS-1-E vs. PPDS-2-E), (C) between 2 and 4 weeks of incubation (PPDS-2-E vs. PPDS-4-E), and (D) 4 and 6 weeks of incubation (PPDS-4-E vs. PPDS-6-E), (E) between 1 and 4 weeks of incubation (PPDS-1-E vs. PPDS-4-E), and (F) 1 and 6 weeks of incubation (PPDS-1-E vs. PPDS-6-E).

The relative amounts of most metabolites were significant higher in embryos than that in megagametophytes at all six time points (Fig. 3). A rather similar pattern of metabolites was found in megagametophyte between successive time points, with a large metabolic change between 0 and 1 WAI (Fig. 4A) and between 2 and 4 WAI (Fig. 4C). A lesser extent of change in metabolites occurred between 1 and 2 WAI (Fig. 4B) and between 4 and 6 WAI (Fig. 4D). After 1 week of incubation, Beta-D-fructose 6-phosphate and ribitol increased 14-fold and decreased 13-fold, respectively (Fig. 4A). In addition, 13 amino acids (L-aspartate, L-glutamine L-threonine, etc.) rapidly increased 1-to 6-fold (Fig. 4A). The changes in phosphorylated sugars and amino acids were relatively little between 2 and 4 WAI (Fig. 4C). However, the relative levels of most fatty acids were declined 1 to 6-fold from 2 WAI to 4 WAI (Fig. 4C). Higher numbers of amino acids and TCA-cycle intermediates were found to be significantly accumulated between 1 and 6 WAI than between 1 and 4 WAI (Fig. 4, E and F).

**Figure 3.**
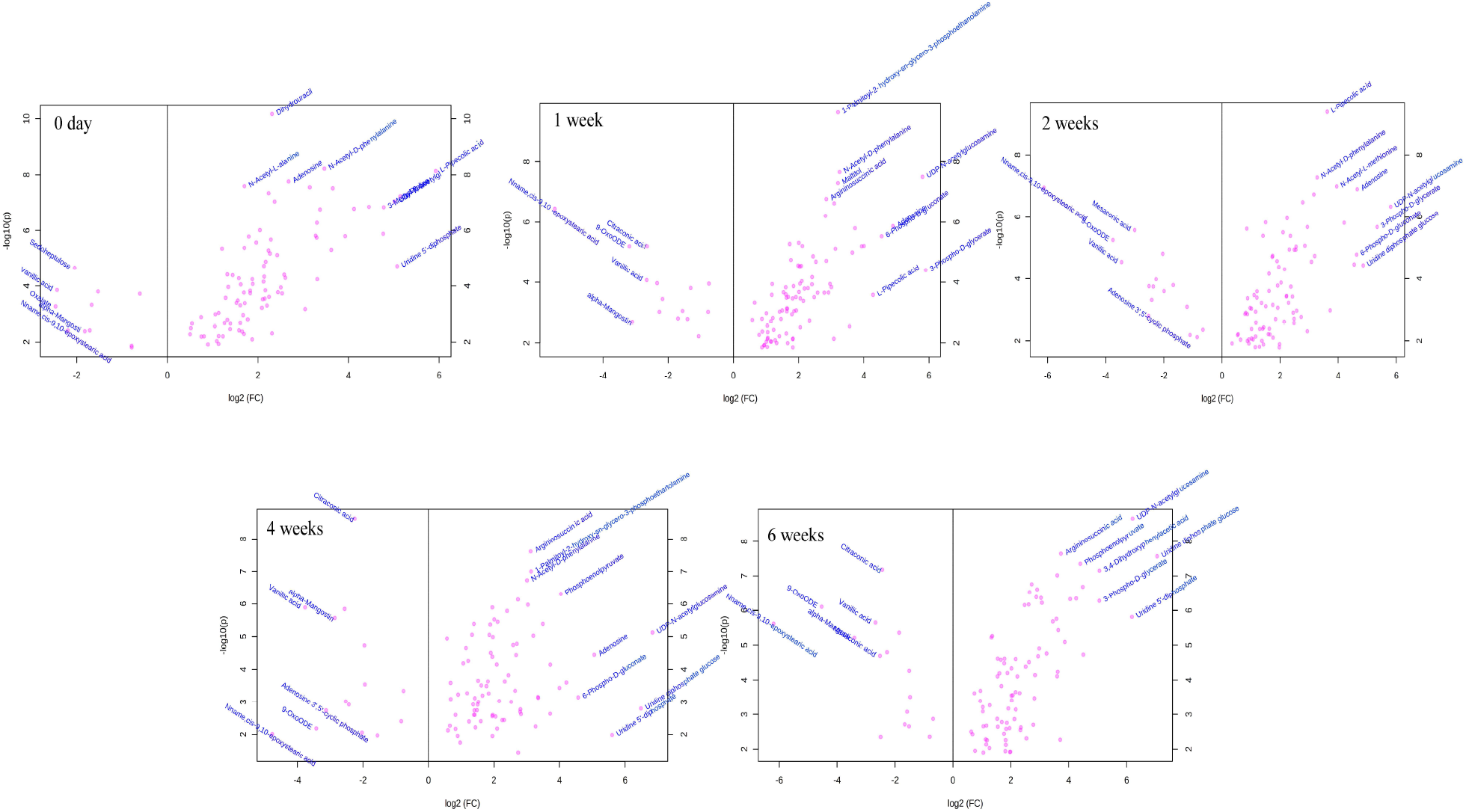
Volcano plots of the important metabolites with VIP value > 1 and P < 0.05 between embryo and megagametophyte of primary physiological dormancy seeds at five different incubation time points.

**Figure 4.**
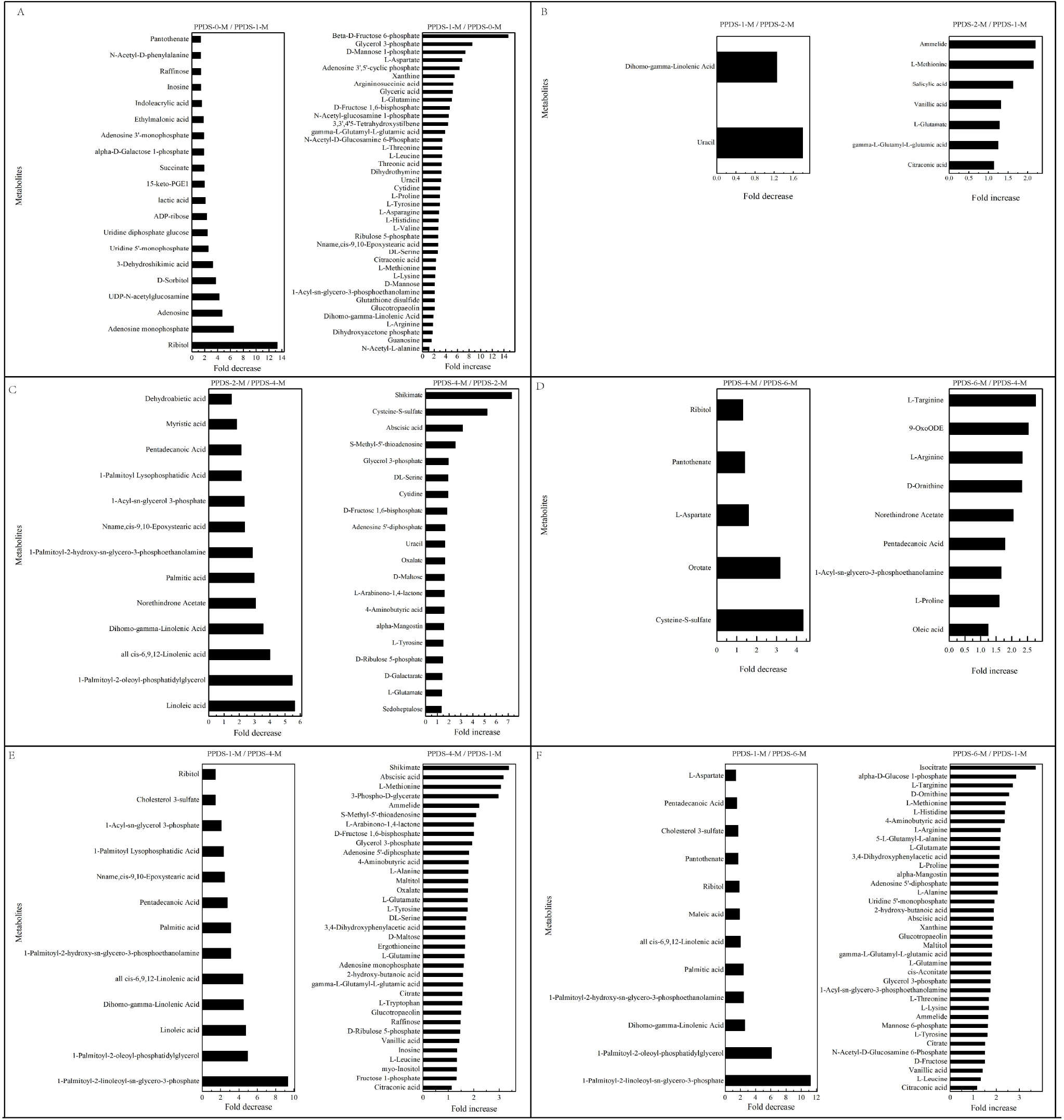
Fold change in the relative contents of important metabolites with VIP value > 1 and *P* < 0.05 in the megagametophytes of primary physiological dormancy seeds (PPDS) (A) between 0 and 1 week of incubation (PPDS-0-M vs. PPDS-1-M), (B) 1 and 2 weeks of incubation (PPDS-1-M vs. PPDS-2-M), (C) between 2 and 4 weeks of incubation (PPDS-2-M vs. PPDS-4-M), (D) 4 and 6 weeks of incubation (PPDS-4-M vs. PPDS-6-M), (E) between 1 and 4 weeks of incubation (PPDS-1-M vs. PPDS-4-M), and (F) 1 and 6 weeks of incubation (PPDS-1-M vs. PPDS-6-M).

### Fold changes of important metabolites with a VIP value > 1 and *P* < 0.05 during incubation of RPPDS

The relative contents of metabolites in the embryos of RPPDS changed little over the first 1 week of incubation (16 increased, 23 decreased) (Fig. 5A). The relative levels of 67 metabolites were significantly reduced in RPPDS with seed coat rupture compared with 1-week-incubated RPPDS (Fig. 5B). In detail, ten phosphorylated sugars (D-mannose 1-phosphate, 6-phospho-D-gluconate, Beta-D-fructose 6-phosphate, alpha-D-glucose 1-phosphate, D-ribose 5-phosphate, D-ribulose 5-phosphate, D-mannose-6-phosphate, L-fucose-1-phosphate, D-fructose 1,6-bisphosphate and fructose 1-phosphate), two sugar alcohols (D-sorbitol and myo-inositol), two disaccharide (D-maltose and trehalose), two trisaccharide (raffinose and stachyose) and 12 amino acids displayed 1-to 8-fold decrease (Fig. 5B). It was found that a surprising 33-fold increase in the relative levels of taxifolin (Fig. 5B). A number of significant changes in metabolite abundance were observed between seed coat rupture and radicle protrusion stages (76 of the 188 metabolites displayed significant changes) (Fig. 5C). There was a 58- and 23 -fold decrease in the respective relative levels of glutathione disulfide and glutathione (Fig. 5C). Beta-D-fructose 6-phosphate, D-mannose 1-phosphate and 6-phospho-D-gluconate dramatically increased 3-to 4-fold (Fig. 5C). However, the differences between the two stages were more subtle compared with the large changes in the metabolome observed from 1 WAI to seed coat rupture stage (Fig. 5, B and C).

**Figure 5.**
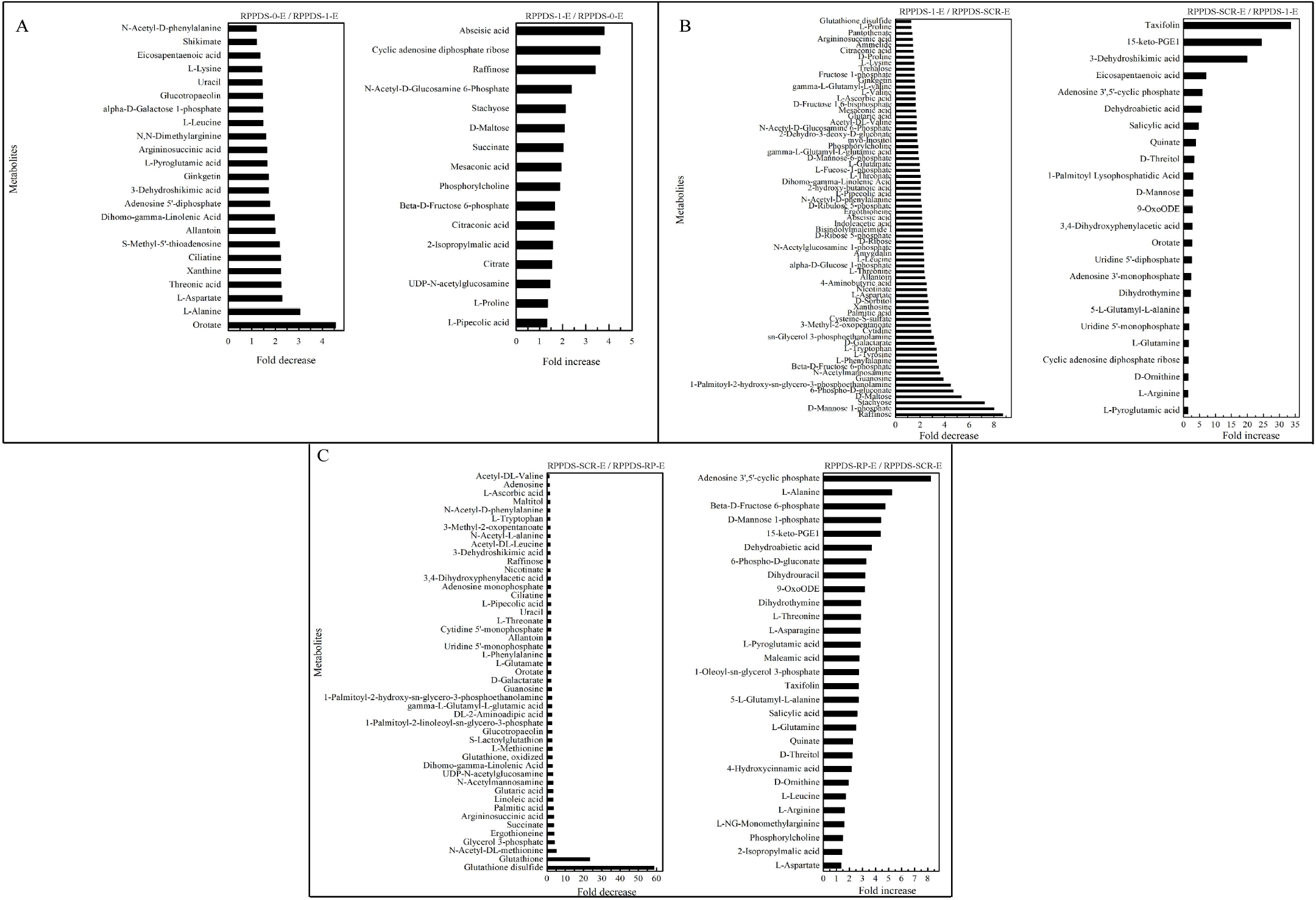
Fold change in the relative contents of important metabolites with VIP value > 1 and *P* < 0.05 in the embryo of seeds with released primary physiological dormancy seeds (RPPDS) (A) between 0 and 1 week of incubation (RPPDS-0-E vs. RPPDS-1-E), and (B) 1 week of incubation and seed coat rupture stage (RPPDS-1-E vs. RPPDS-SCR-E), and (C) seed coat rupture stage and radicle protrusion stage (RPPDS-SCR-E vs. RPPDS-RP-E).

At 0 WAI, 1 WAI and seed coat rupture stage, the embryos in RPPDS contained higher relative contents of most metabolites than megagametophytes (Fig. 6). At 1 WAI, a smaller proportion of the detected metabolites with VIP > 1 and P < 0.05 changed in megagametophytes (Fig. 7A). The trend of changes in metabolites was then followed by the largest change between 1 WAI and seed coat rupture stage (Fig. 7B), followed by a relatively small change between seed coat rupture and radicle protrusion stages (Fig. 7C). During the period form 1 WAI to seed coat rupture stage, two sugar alcohols (maltitol and myo-inositol), one disaccharide (D-maltose), and three trisaccharide (raffinose, stachyose and maltotriose) significantly decreased 1-to 5-fold (Fig. 7B). While most of the changes in amino acids were seen to occur in megagametophyte, 15 amino acids displaying 1.4- and 2.7-fold increases at seed coat rupture stage (Fig. 7B).

**Figure 6.**
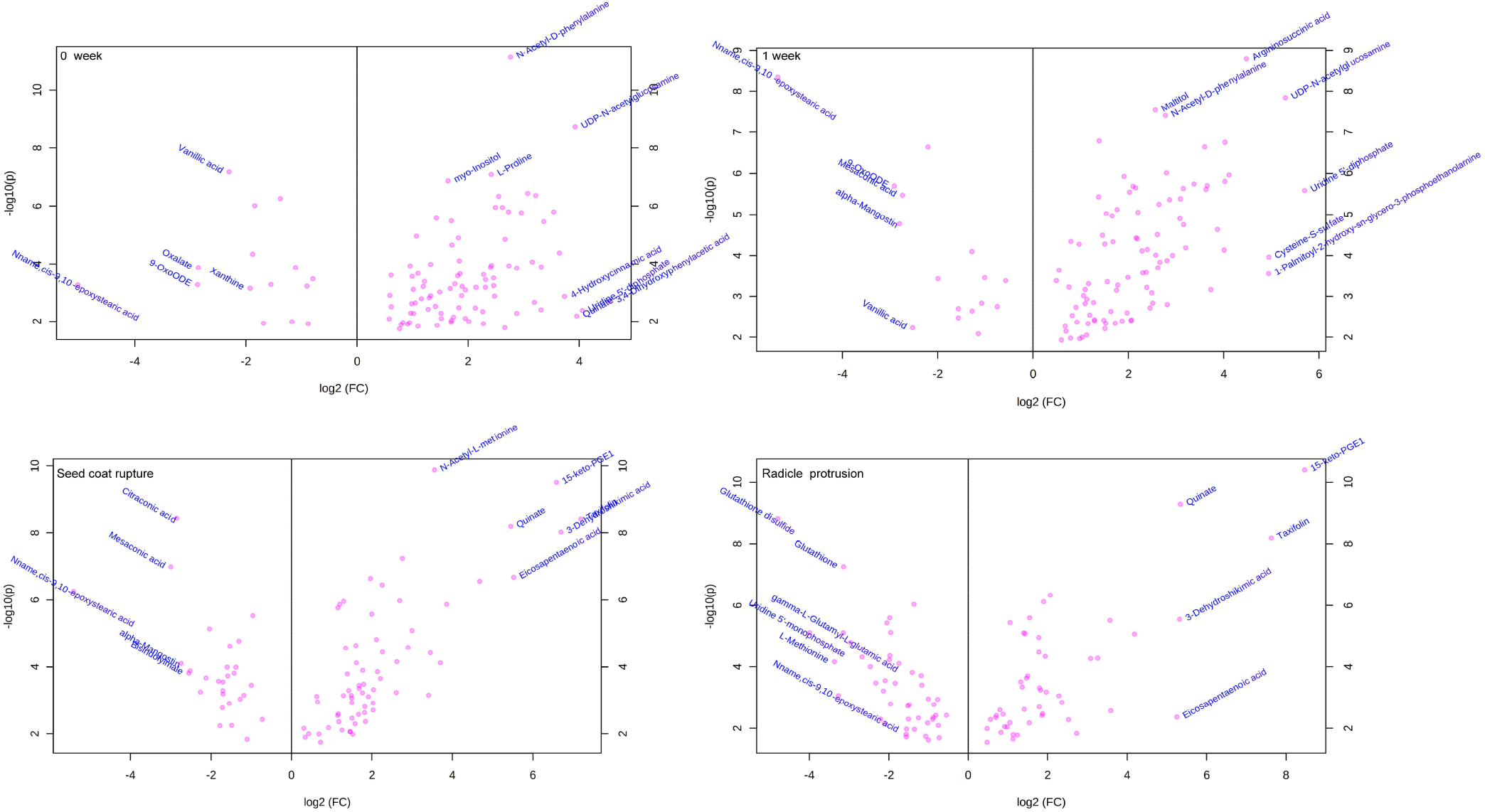
Volcano plots of the important metabolites with VIP value > 1 and P < 0.05 between embryo and megagametophyte of seeds with release primary physiological dormancy at four incubation time points.

**Figure 7.**
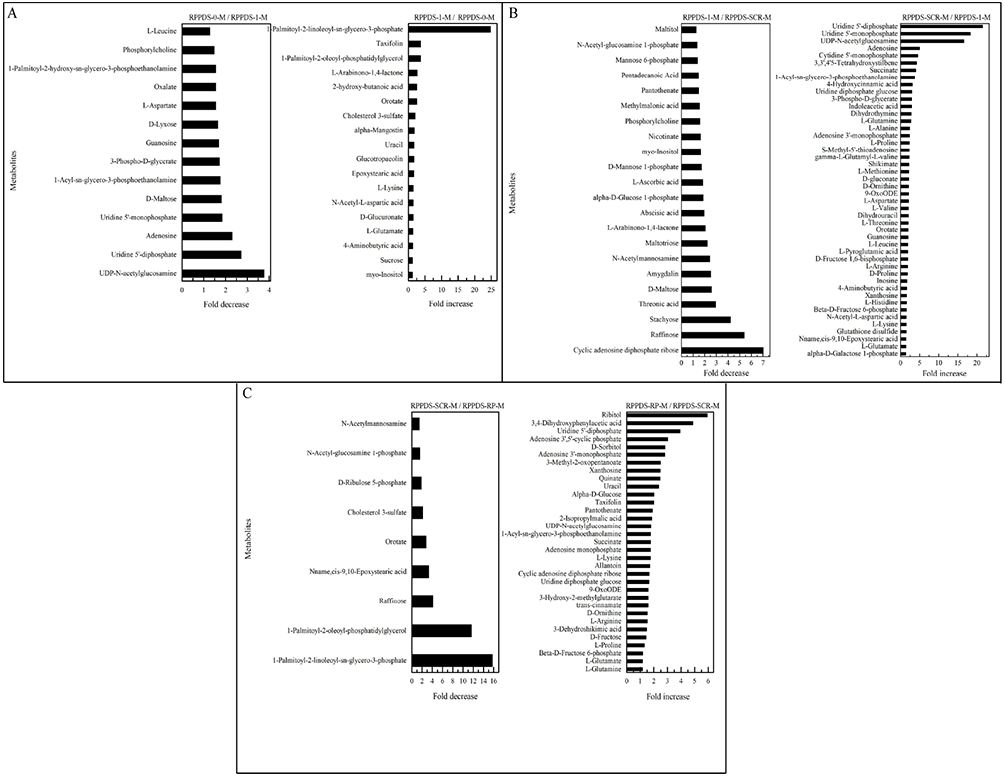
Fold change in the relative contents of important metabolites with VIP value > 1 and *P* < 0.05 in the megagametophyte of seeds with released primary physiological dormancy seeds (RPPDS) (A) between 0 and 1 week of incubation (RPPDS-0-M vs. RPPDS-1-M), and (B) 1 week of incubation and seed coat rupture stage (RPPDS-1-M vs. RPPDS-SCR-M), and (C) seed coat rupture stage and radicle protrusion stage (RPPDS-SCR-M vs. RPPDS-RP-M).

### Fold changes of important metabolites with a VIP value > 1 and *P* < 0.05 between PPDS-4W and RPPDS-1W, PPDS-4W and RPPDS-SCR

The metabolic process occurred before radicle protrusion regulate the complete of germination. In addition, the large metabolic changes in PPDS were observed after 4 weeks of incubation. Thus, we performed the fold change analyses of metabolites differentially accumulated between 1-week-incubated RPPDS, RPPDS with seed coat rupture and 4-weeks-incubated PPDS, aiming at reveal real potential metabolic processes that control seed primary physiological dormancy maintenance. 92 metabolites were found to be significantly accumulated in the embryos between PPDS-4W and RPPDS-1W (Fig. 8A). Of these, the relative levels of 41 metabolites were significant higher in PPDS-4W, including 6-phospho-D-gluconate, abscisic acid, raffinose, stachyose, alpha-D-glucose 1,6-bisphosphate, Beta-D-fructose 6-phosphate, D-ribulose 5-phosphate, D-mannose 1-phosphate, D-fructose 1,6-bisphosphate and citrate, etc. The relative levels of most amino acids were higher in RPPDS-1W than that in PPDS-4W. Only the relative levels of 27 metabolites were significantly different in embryos between RPPDS-SCR and PPDS-4W (Fig. 8B). 6-phospho-D-gluconate, D-mannose 1-phosphate and Beta-D-fructose 6-phosphate accumulated 10-to 48-fold in PPDS-4W. However, taxifolin and 3-dehydroshikimic acid decreased 62- and 86-fold, respectively, in PPDS-4W compared to RPPDS-SCR.

**Figure 8.**
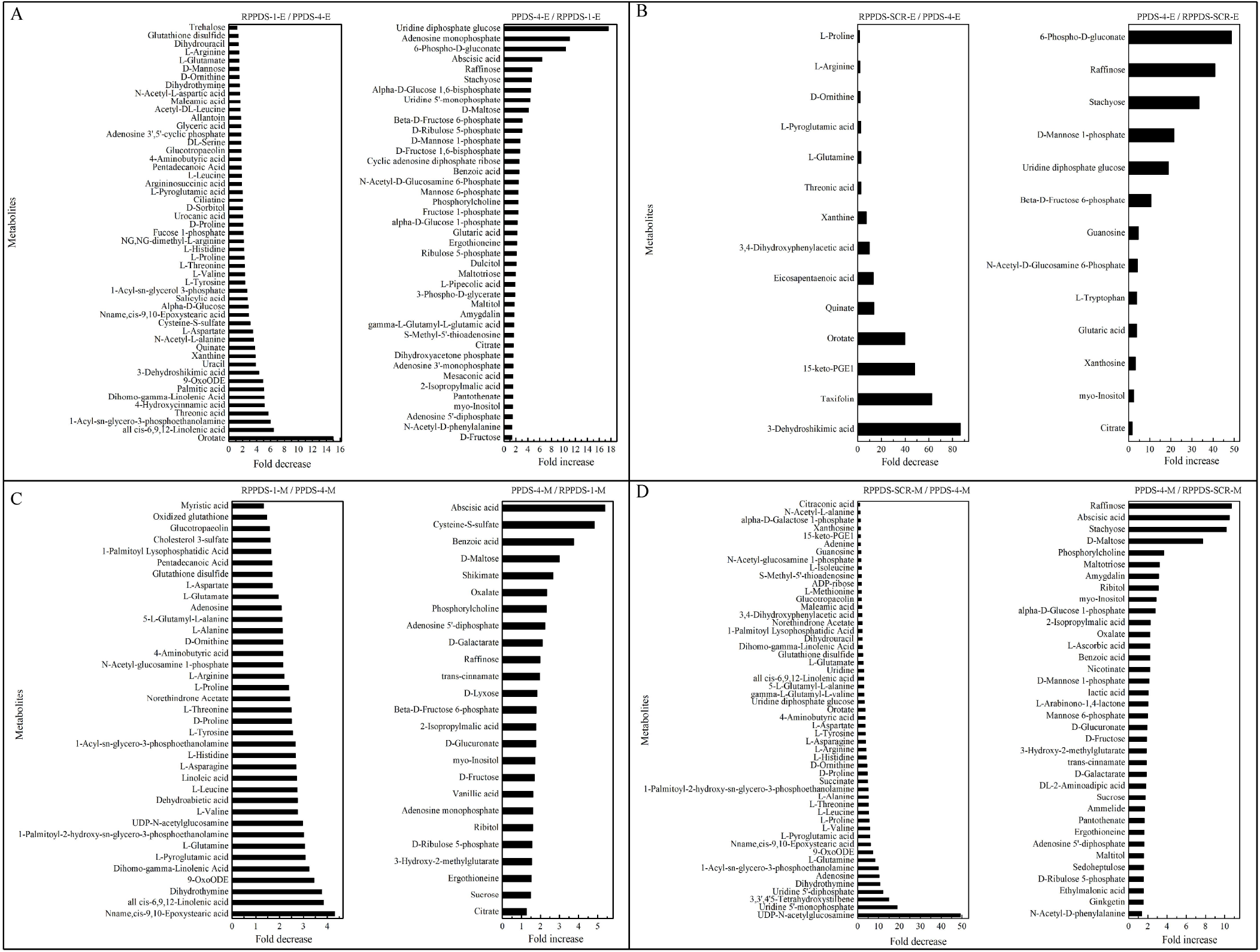
Fold change in the relative contents of important metabolites with VIP value > 1 and *P* < 0.05 in the embryo of (A) between seeds with released primary physiological dormancy seeds (RPPDS) incubated for 1 week (RPPDS-1-E) and primary physiological dormancy seeds (PPDS) incubated for 4 weeks (PPDS-4-E), (B) between RPPDS with seed coat rupture (RPPDS-SCR-E) and PPDS-4-E, in the megagametophytes of (C) between seeds with released primary physiological dormancy seeds (RPPDS) incubated for 1 week (RPPDS-1-M) and primary physiological dormancy seeds (PPDS) incubated for 4 weeks (PPDS-4-M), and (D) between RPPDS with seed coat rupture (RPPDS-SCR-M) and PPDS-4-M.

The relative content of abscisic acid in megagametophyte was 5.3-fold higher in PPDS-4W rather than in RPPDS-1W (Fig. 8C). D-maltose, raffinose, D-lyxose, myo-inositol, D-fructose, ribitol and sucrose also remained high relative levels in PPDS-4W, whereas 14 amino acids were at least twofold more highly accumulated in RPPDS-1W compared with PPDS-4W. More amino acids were remained in the megagametophytes of RPPDS-SCR than in PPDS-4W (Fig. 8D). In contrast, raffinose, stachyose, D-maltose, maltotriose, ribitol, myo-inositol, D-fructose, sucrose, maltitol and sedoheptulose exhibited higher relative contents in PPDS-4W.

### The patterns of change in metabolic pathways during incubation of PPDS and RPPDS

Most metabolites whose relative levels significantly changed during incubation of PPDS or RPPDS were mainly associated with carbohydrate metabolism, amino acid metabolism and lipid metabolism (Table 1 and 2). The activities of the above-mentioned most metabolic processes were higher in embryos than that in megagametophytes at the various time points of incubation. For PPDS, a dramatic change in most metabolic pathways was observed at both 1 WAI and 4 WAI (Table 1). There was also a strong metabolic switch from 1 WAI to seed coat rupture stage in RPPDS (Table 2).

**Table 1.**
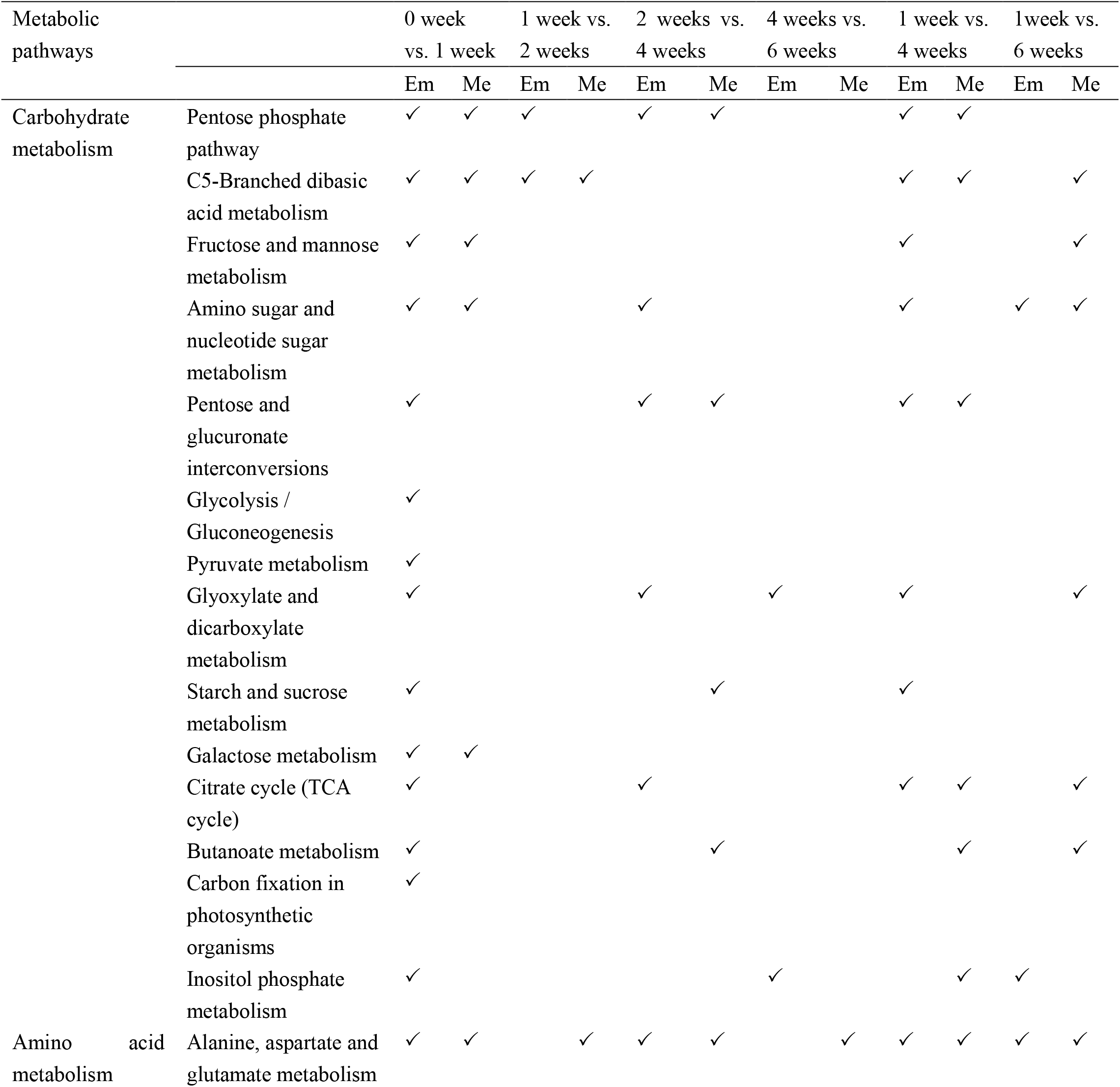
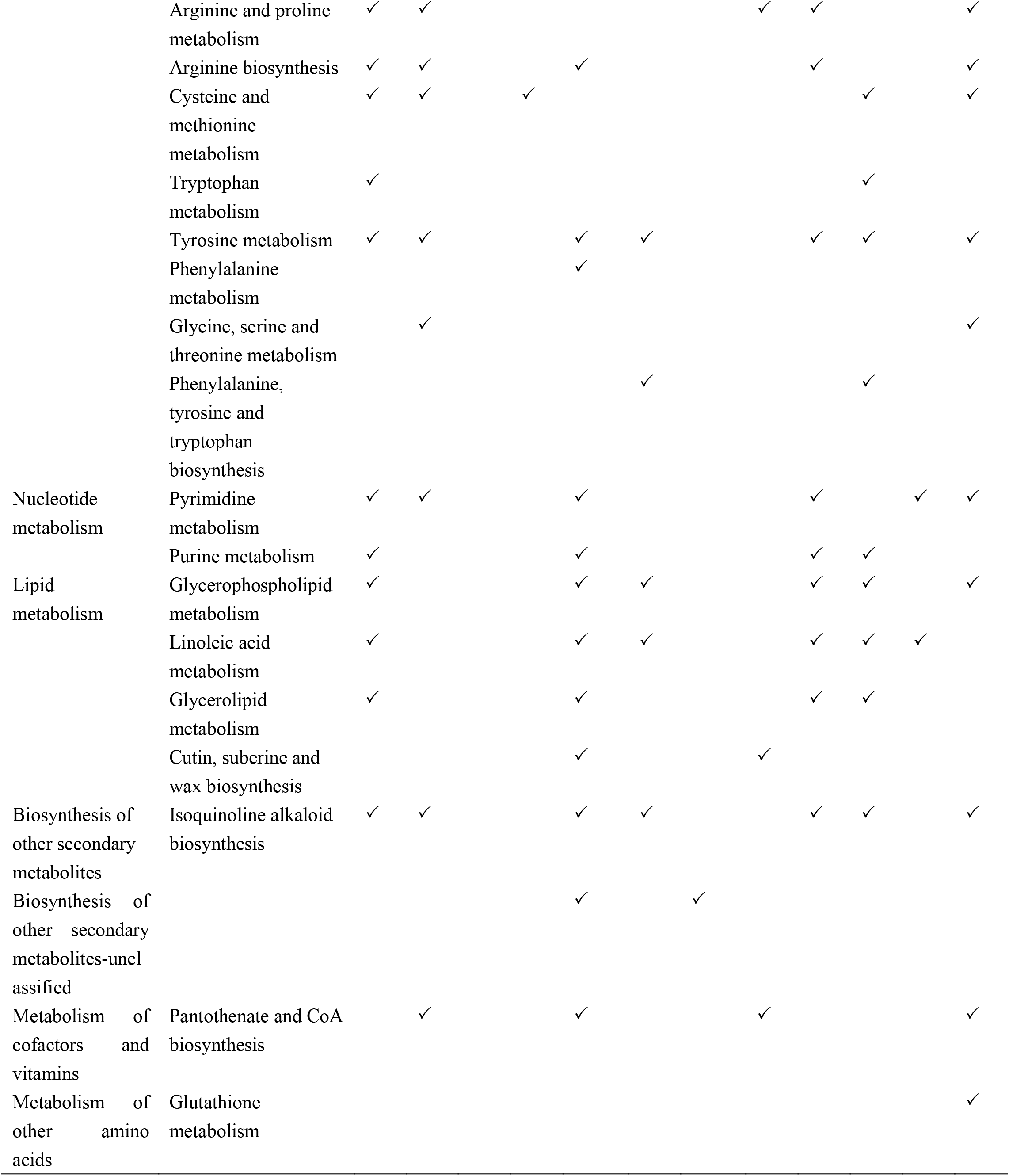
The altered metabolic pathways in the embryo and megagametophyte of primary physiological dormancy seeds between different incubation time points. Em: embryo, Me: megagametophyte. “✓” indicates significantly changed metabolic pathway between two time points.

**Table 2.**
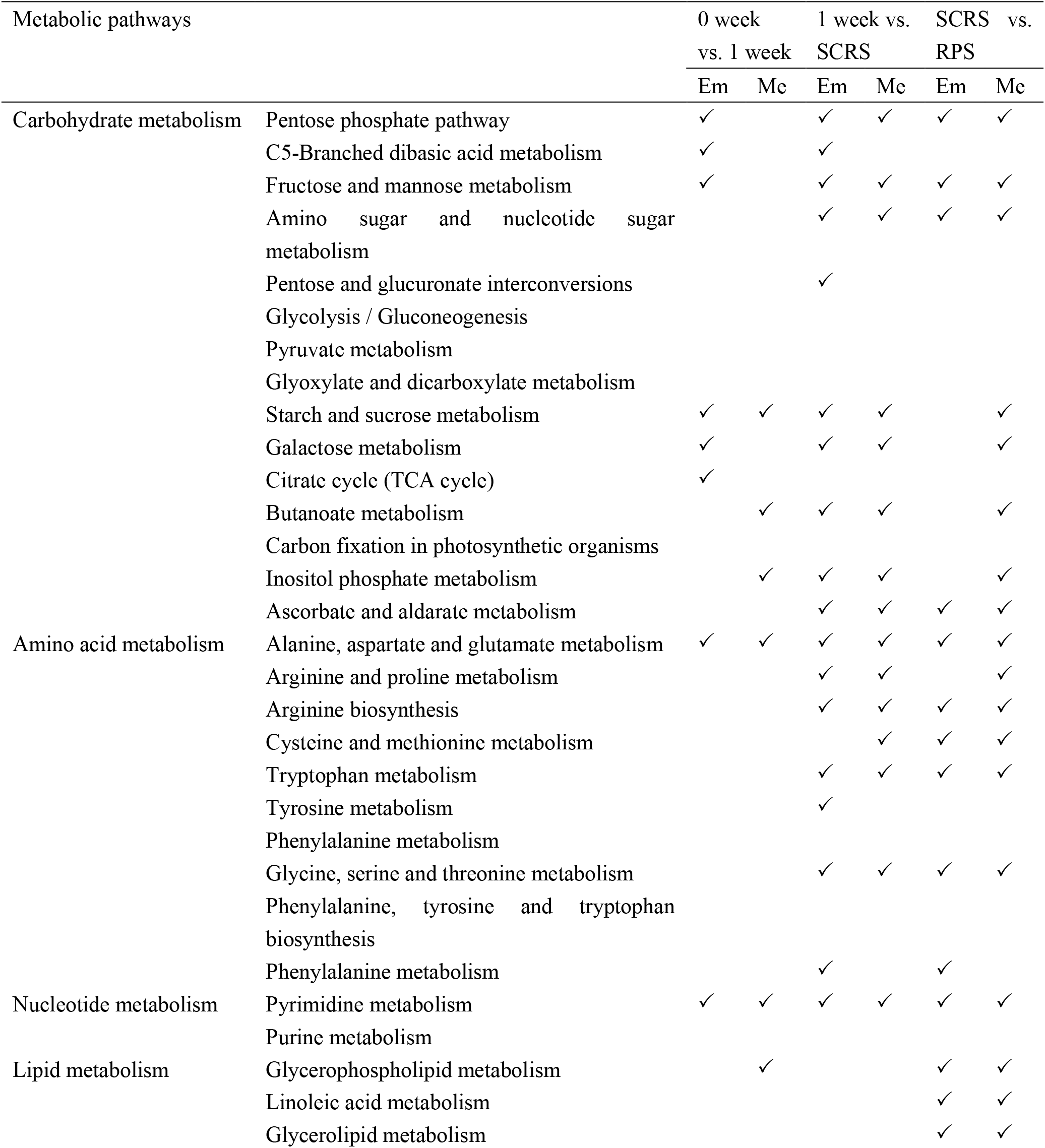
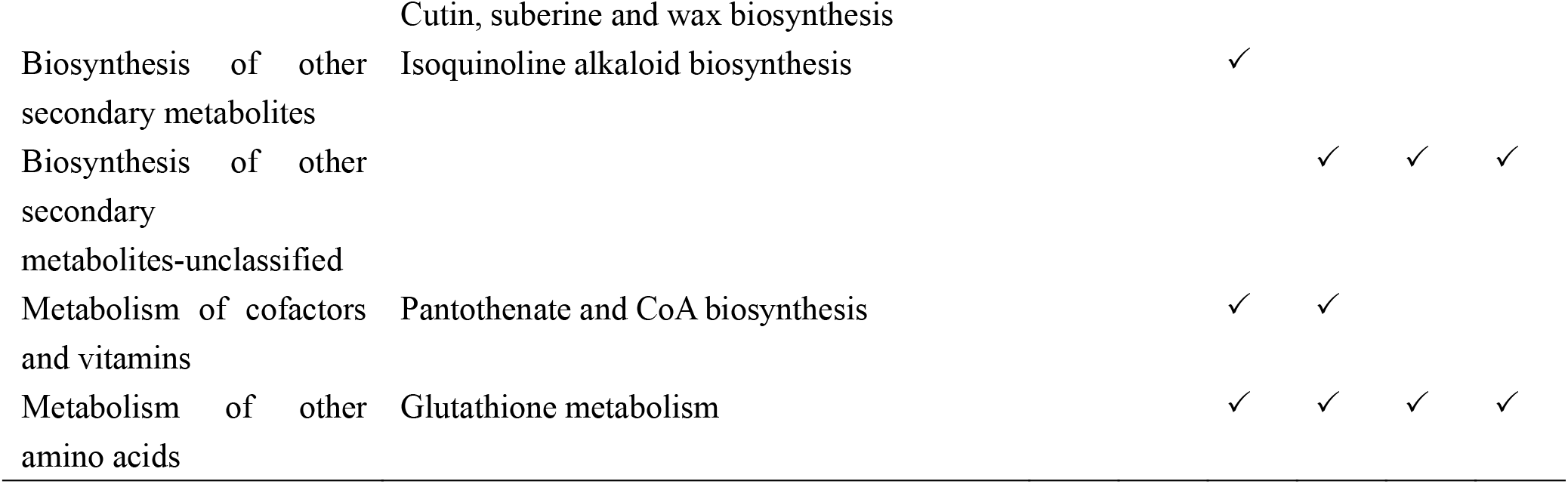
The altered metabolic pathways in the embryo and megagametophyte of seeds with released primary physiological dormancy between successive time points. Em: embryo, Me: megagametophyte. SCRS: seed coat rupture stage, RPS: radicle protrusion stage. “✓” indicates significantly changed metabolic pathway between two time points.

### Differential metabolic pathways between PPDS-4 and RPPDS-1, between PPDS-4 and RPPDS-SCR

Carbohydrate metabolism and amino acid metabolism were the mainly changed metabolic pathways between PPDS-4 and RPPDS-1, between PPDS-4 and RPPDS-SCR (Table 3). There were more differential metabolic pathways in embryo between PPDS-4 and RPPDS-1 than between PPDS-4 and RPPDS-SCR.

**Table 3.**
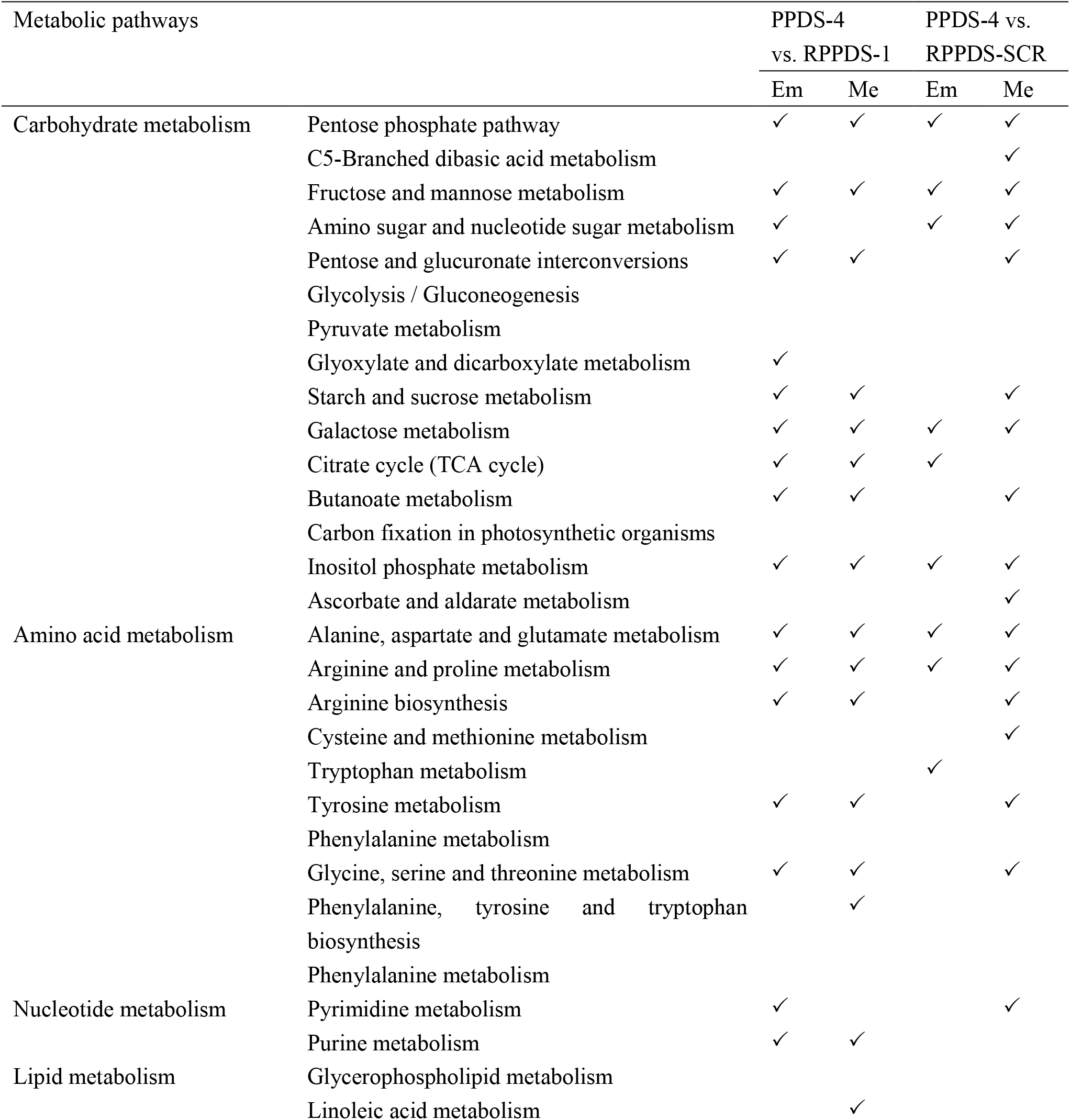
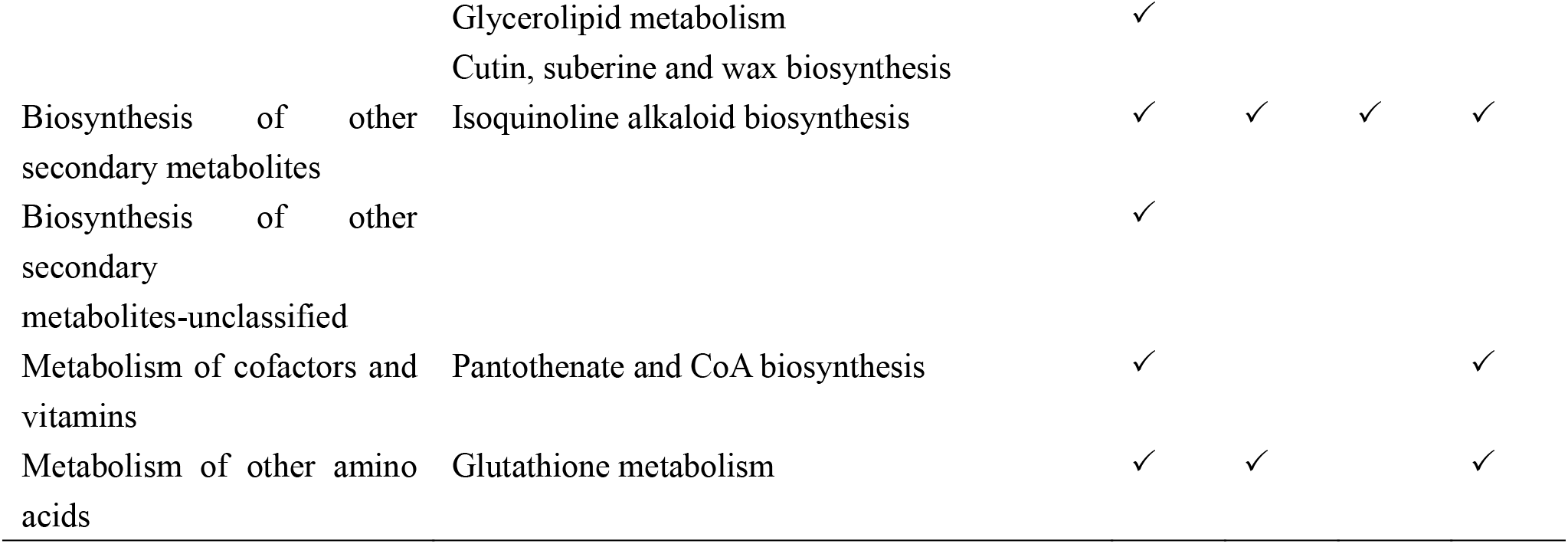
The altered metabolic pathways in the embryo and megagametophyte between 4-weeks-incuabted primary physiological dormant seeds (PPDS-4) and 1-weeks-incubated seeds with released primary physiological dormancy (RPPDS-1), between PPDS-4 and seeds with released primary physiological dormancy at seed coat rupture stage (RPPDS-SCR). Em: embryo, Me: megagametophyte. “✓” indicates significantly changed metabolic pathway.

## DISCUSSION

This study provides a comprehensive profile of the metabolites during germination of Korean pine seeds. After dry PPDS were imbibed for 1 week under favorable conditions, most metabolites in embryos with significant FC and VIP > 1 were involved in carbohydrate metabolism and amino acid metabolism. In detail, there was a great increase in the relative levels of 6-phospho-D-gluconate, D-mannose 1-phosphate, 3-phospho-D-glycerate, Beta-D-fructose 6-phosphate, phosphoenolpyruvate, ribulose 5-phosphate, D-fructose 1,6-bisphosphate and fructose 1-phosphate associated with pentose phosphate pathway, fructose and mannose metabolism and glycolysis. This in accordance with our previous studies that showed the glycolytic pathway activated in the dormant Korean pine seeds upon imbibition (Song and Zhu, 2019). It has been reported that a rapid uptake of water occurred during first 5 days of incubation of dry dormant Korean pine seeds. The increase in various metabolic activities during this period might be caused by water uptake. Further investigations about the metabolic changes that occur in dormant seeds with the extension of incubation time are thus needed to distinguish the consequences of water absorption from the metabolic activities involved in seed dormancy maintenance. We found that these early responses to water were then followed by relatively small changes in the relative contents of metabolites between 1 and 2 WAI. In contrast, there was a larger metabolic switch occurs as the incubation time was extended to 4 weeks, with the relative levels of 45 of the 59 metabolites with significant FC and VIP > 1 were increased between weeks2 and 4 of incubation. Furthermore, both PPP and TCA ran at a higher rate at the 4 WAI time point when compared to 1 WAI. In addition, glycolysis-cycle-related metabolites, D-fructose 1,6-bisphosphate and alpha-D-glucose 1-phosphate, were also accumulated. However, this accumulation did not last until the 6 WAI time point. A transient but rapid increase in most metabolites occurred after 4 weeks of incubation when dormant seeds had already been imbibed. Based on these findings, we proposed that dormant Korean pine seeds are far from metabolically quiescent state when they were in favorable conditions for germination of non-dormant seeds. Cadman et al. (2006) also found that transcriptional activities took place in primary dormant (PD) Arabidopsis Cvi seeds after 30 days of imbibition due to 1833 genes were highly expressed compared with 24-h-imbibed PD seeds.

Only a small proportion of the metabolites such as L-alanine, L-aspartate and N-(L-arginino) succinate, were differentially accumulated in the embryos of RPPDS during the first 1 week of incubation. However, the relative levels of those metabolites involved in the carbohydrate metabolic pathways including fructose and mannose metabolism, TCA cycle, PPP, galactose metabolism and starch and sucrose metabolism were significantly reduced. The pattern of change in metabolic activities described above is contrary to our earlier statements that Korean pine seeds rapidly resumed metabolic activity upon incubation in suitable conditions for germination (Song and Zhu, 2019). In the present study, when RPPDS were retrieved from virgin MKPF forest in late May, these seeds had previously experienced the relatively high temperature conditions in soil, which might be the reason for the slow increase in the relative levels of the most metabolites under laboratory conditions. That is, a great metabolites increase was not observed in the present study because of the delay in sampling time. In the embryos of RPPDS with seed coat rupture, there was a major decrease in the relative levels of various phosphorylated sugars (such as D-mannose 1-phosphate, 6-phospho-D-gluconate, Beta-D-fructose 6-phosphate, alpha-D-glucose 1-phosphate, D-ribulose 5-phosphate, etc.), polysaccharides (D-maltose and trehalose), oligosaccharides (raffinose and stachyose) and most amino acids when compared to 1-week-incubated RPPDS. Based on these results we hypothesize that the metabolic activity is attenuated in the embryos of RPPDS with seed coat rupture. Moreover, TCA-cycle was also significantly down-regulated relative to RPPDS. Non-Targeted metabolomics analysis of Poplar seeds also revealed that three TCA cycle intermediates (citrate, isocitrate and succinate) substantially accumulated during the periods of slow water uptake and then promptly decreased until radicle protrusion (Qu et al., 2019). Research conducted with non-dormant and dormant seeds of *Leymus chinensis* demonstrated that the proteins involved in TCA cycle were decreased in non-dormant seeds (Hou et al., 2009). When the radicle protruded through the seed coat, part of the embryo was in an atmosphere containing sufficient oxygen, PPP was significantly enhanced, while TCA cycle did not exhibit dramatic change. Allen et al. (2010) suggested that fermentation still proceeded in Arabidopsis seeds approaching radicle protrusion.

It is generally accepted that glycolysis, fermentation, the TCA cycle and PPP are activated to provide energy for metabolic activity during seed germination of non-dormant seeds (Fait et al., 2006; Preston et al., 2009; Angelovici et al., 2011; Weitbrecht et al., 2011; He and Yang, 2013; Zaynab et al., 2018). It has been also suggested that because of limited oxygen or lack of functional mitochondria, energy is mainly provided by glycolysis and fermentation during early stage of germination (Yang et al., 2007; He et al., 2011b; He and Yang, 2013; Ren et al., 2018; Zaynab et al., 2018), and TCA cycle is the main energy source at the late stage of germination. A decrease in TCA cycle activity would lead to the inhibition of germination of dormant seeds (Chibani et al., 2006; Angelovici et al., 2011; Das et al., 2017; Ren et al., 2018). However, we found that the relative levels of a number of metabolites related to carbohydrate metabolism were significant higher in the embryos of 4-weeks-incubated PPDS, especially those related to PPP and TCA cycle, when compared to 1-week-incubated RPPDS. Xia et al. (2018a) also detected a higher activity of enzymes related to TCA cycle and glycolysis in imbibed-dormant sunflower seeds. Furthermore, sunflower seed dormancy can be released by cyanide (a respiratory inhibitor), implying the importance of respiratory activity in seed dormancy maintenance (Oracz et al., 2008; 2009).

Conversion of glucose through glycolysis, β-oxidation and TCA cycle yields nicotinamide adenine dinucleotide (NADH) and reduced flavin adenine dinucleotide (FADH_2_) which serve as electron donors for the generation of adenosine triphosphate (ATP) in the mitochondrial electron transport chain (Liemburg-Apers et al., 2015). The electron escaped from electron transport chain (ETC) can lead to the generation of reactive oxygen species (ROS) (Liemburg-Apers et al., 2015). NADPH-oxidases are also involved in the production of ROS (Ye et al., 2012). Although it has been documented that ROS accumulation in the cells might damage proteins and membranes, ROS also act as a key positive regulatory factor in the activation of germination process (Murthy et al. 2003; Bailly, 2004; Oracz et al., 2007; El-Maarouf-Bouteau and Bailly, 2008; Müller et al., 2009; Parkhey et al. 2012; Ye et al., 2012). Sugar enters to glycolysis and further TCA cycle mainly through phosphorylation of sugars (Fait et al., 2006). A significant increase of numerous phosphorylated sugars and organic acids involved in glycolysis, PPP and TCA cycle during incubation of PPDS may indicate an overaccumulation of ROS, leading to the maintenance of seed dormancy. There was a 1-and 10-fold increase in the respective levels of adenosine triphosphate (ADP) and adenosine monophosphate (AMP) in 4-weeks-incubated PPDS compared with 1-week-incubated RPPDS. In addition, the ratio of glutathione to glutathione disulfide was also slightly higher in 4-weeks-incubated PPDS. These results suggested that the PPDS might be subjected to more severe oxidative stress condition. In contrast, the activities of TCA cycle and PPP are attenuated most presumably to reduce the generation of ROS during incubation of RPPDS under favorable conditions for germination. The reduced TCA cycle and PPP may reflect a way to avoid energy waste and reserve consumption, preparing for the establishment of autotrophic growth of seedling after seed germination.

It has been confirmed that abscisic acid (ABA) plays a critical role in the induction and expression of seed dormancy and the inhibition of seed germination (Ali-Rachedi et al., 2004; Liu et al., 2013). Xia et al. (2018b) reported that the ABA level significantly accumulated in dormant sunflower seeds, but continued to decrease in non-dormant seeds after 24 h of imbibition. The 24 h imbibition was also the time when the metabolism of non-dormant seeds was clearly differentiated from that of dormant seeds. A rather similar pattern of change in the relative levels of ABA was found in the present study, with a great increase in embryos of PPDS after 4 weeks of incubation and a rapid decline in embryos of RPPDS after 1 week of incubation. Similarly, there was a great variation in metabolism between the 4-weeks-incubated PPDS and 1-week-RPPDS. It is therefore to hypothesize that ABA may affect seed dormancy (or seed germination) process by its action on metabolic actives. Metabolic, transcriptomic and proteomic analyses on seeds treated with exogenous ABA also showed that the regulation of ABA in seed germination is involved in the inhibition of some metabolic processes (Chibani et al. 2006; Xia et al., 2018b).

In the present study, all measured metabolic pathways were more active in embryo than that in megagametophyte in both PPDS and RPPDS. Carbohydrate metabolism changed similarly to amino acid metabolism in embryo during incubation. However, in megagametophyte, amino acid metabolism was altered much more prominently than carbohydrate metabolism. In addition, there was a more dramatic change in amino acid metabolism in megagametophytes between 4-weeks incubated PPDS and 1-week-incubated RPPDS. It appears that both carbohydrate metabolism and amino acid metabolism are more predominant metabolic pathways in embryo, but in megagametophyte amino acid metabolism become dominant. Similarly, several studies have demonstrated that the embryos mainly contain enzymes that related to ATP synthesis, energy generation or citrate synthesis (Penfield et al., 2006; Galland et al., 2017; Liu et al., 2018; Domergue et al., 2019). In contrast, catabolic enzymes were found in the endosperm. Xu et al. (2016) reported that proteins with increased abundance in embryo were mainly associated with glycolysis and TCA cycle, cell growth and division and protein synthesis during dormancy release of wild rice seeds. While, the abundances of these proteins in endosperms maintained steady-state level or decreased.

The relative levels of amino acids in either embryos or megagametophytes were always significant higher in RPPDS than that in PPDS during incubation. A great decrease in most amino acids occurred from 1 week after incubation to seed coat rupture in the embryos of RPPDS, suggesting that amino acids might be utilized for protein biosynthesis required for seed germination. In contrast, only a few amino acids exhibited a slow increasing trend in embryos during incubation of PPDS. Thus, we concluded that the attenuated amino acid metabolism may maintain primary physiological dormancy in Korean pine seeds. Previous studies have shown that protein biosynthesis was inhibited in dormant seeds compared with non-dormant seeds (Bove etal., 2005; Cadman et al. 2006; Carrera et al., 2008).

## MATERIALS AND METHODS

### Seed collection

In late September 2018, fresh Korean pine seeds were collected from virgin MKPF in Wuying fenglin National Nature Reserves in northeastern China (128°58′–129°15′E, 48°02′– 48°12′N). Seeds contained a low amount of water (about 10%). These fresh dry seeds were then stored at −20 °C to maintain their primary physiological dormancy status.

### Seed burial experiment

In 20 October 2018, a proportion of Korean pine seeds were buried between litterfall and soil in virgin MKPF to release primary physiological dormancy under the effect of lower late autumn and winter temperatures in natural conditions. At the same time, another proportion of seeds were continually kept at −20°C. These seeds were considered as primary physiological dormancy (PPDS).

### Incubation experiment

In late May 2019, seeds were retrieved from virgin MKPF and considered as seeds with released primary physiological dormancy (RPPDS). Both PDS and RPPDS were used for incubation experiment. Seeds were incubated in a growth incubator under a temperature fluctuation regime of 25°C for 14h and 15°C for 10h. An approximately 200 μmol of photons m^-2^ s^-1^ was provided by cool white fluorescent tubes during high temperature (14h) and darkness was set during low temperature (10h). Group of 20 PPDS (or RPPDS) were placed in a 10-cm Petri dish with 5 layers of filter paper moistened with 10 ml deionized water. Petri dish was then covered with parafilm to reduce water loss. PPDS were incubated for six weeks and sampled on week 0, week 1, week 2, week 4 and week 6 after incubation. The PPDS collected at five time points were abbreviated to PPDS-0, PPDS-1, PPDS-2, PPDS-4 and PPDS-6, respectively. RPPDS were incubated and sampled at four stages (0 week of incubation, 1 week of incubation, seed coat rupture stage and radicle protrusion stage). The RPPDS collected at four stages were abbreviated to RPPDS-0W, RPPDS-1W, RPPDS-SCR and RPPDS-RP, respectively. At each sample time point, five Petri dishes were taken out from growth incubator and used as five replicates.

### Seed samples pretreatment

The relative levels of metabolites in the embryos of five PPDS samples (PPDS-0-E, PPDS-1-E, PPDS-2-E, PPDS-4-E and PPDS-6-E), the megagametophytes of five PPDS samples (PPDS-0-M, PPDS-1-M, PPDS-2-M, PPDS-4-M and PPDS-6-M), the embryos of four RPPDS samples (RPPDS-0-E, RPPDS-1-E, RPPDS-SCR-E and RPPDS-RP-E) and the megagametophytes of four RPPDS samples (RPPDS-0-M, RPPDS-1-M, RPPDS-SCR-M and RPPDS-RP-M) were all used for a non-targeted metabolome analysis. The excised embryos (or megagametophytes) were immediately frozen in liquid nitrogen and subsequently stored at −80°C. Embryos (or megagametophytes) samples were ground into powder under the protection of liquid nitrogen. 80 mg embryo (or megagametophyte) powders were placed inside an Eppendorf tube. One ml of precooled extraction solution (methanol/acetonitrile/ddH_2_O (2:2:1, v/v/v)) was then added. The mixture of powder and extraction solution was then subjected for following treatments, vortex for 60 seconds, low-temperature ultrasonic for 30 minutes (twice), stand at −20°C for 1 hour to precipitate protein, centrifugation for 20 minutes (14,000 RCF, 4°C). After that, the supernatant was lyophilized and stored at −80°C until used for further metabolomics analysis.

### LC-MS analysis

The UHPLC/MC system consisted of an Agilent 1290 Infinity LC (Agilent Technologies, Santa Clara, CA, USA) connected to a TripleTOF 6600 mass spectrometer (AB SCIEX, USA). Chromatographic analyses were performed using a hydrop interaction liquid chromatography (HILIC) column (ACQUITY UPLC BEH Amide 2.1 mm × 100 mm column, internal diameter 1.7 μm, Waters, Ireland) maintained at 25°C, and the autosampler tray temperature was maintained at 4 °C. Mobile phase A was water with 25 mM ammonium acetate and 25 mM ammonia; mobile phase B was acetonitrile. The gradient conditions were 0-1 min, 95% B; 1-14 min, from 95 to 65% B; 14-16 min, from 65 to 40% B; 16-18 min, 40% B; 18-18.1 min, from 40 to 95% B and 18.1-23 min, 95% B. The flow rate was 0.3 ml minute^-1^, and the injection volume was 2μL. For both modes of operation, the mass spectrometric parameters were like that used in Zhou et al. (2019).

## Data analysis

Raw data (wiff. scan files) were converted to mzXML format using ProteoWizard. Then, peak alignment, retention time correction, and peak area extraction was conducted with XCMS program.

Metabolite structure identification was performed with accurate mass number matching (< 25ppm) and secondary spectrum matching. The ion peaks with greater missing values (>50%) were removed. Metabolites were searched in a laboratory self-built database. Statistical analyses were conducted with Metaboanalyst (www.metaboanalyst.ca/).

Principal component analysis (PCA) was used to globally investigate the metabolic changes in both embryo and megagametophyte during incubation of PPDS (or RPPDS). For PCA analysis performed on the relative contents of metabolites in the embryos and megagametophytes of five types of PPDS, the metabolite data were log transformed (generalized logarithm transformation) and auto scaled (mean-centered and divided by the standard deviation of each variable) for normalization. To gain a better normalization result, pareto scaling (mean-centered and divided by the square root of the standard deviation of each variable) was only used for the metabolite data obtained from the embryos and megagametophytes of four types of RPPDS. We also used PCA to analyze the alterations in metabolic profiles of the embryos (or megagametophytes) of both PPDS and RPPDS. The metabolite data of the embryos of PPDS and RPPDS were log transformed and auto scaled for normalization, and then used for PCA analysis. The metabolite data of the megagametophytes of PPDS and RPPDS were also subjected to PCA analysis after auto scaled for normalization.

Partial least squares-discriminant analysis (PLS-DA) was conducted over 20 pairs of samples (PPDS-0-E vs. PPDS-1-E, PPDS-1-E vs. PPDS-2-E, PPDS-2-E vs. PPDS-4-E, PPDS-4-E vs. PPDS-6-E, PPDS-1-E vs. PPDS-4-E, PPDS-1-E vs. PPDS-6-E, PPDS-0-M vs. PPDS-1-M, PPDS-1-M vs. PPDS-2-M, PPDS-2-M vs. PPDS-4-M, PPDS-4-M vs. PPDS-6-M, PPDS-1-M vs. PPDS-4-M, PPDS-1-M vs. PPDS-6-M, RPPDS-0-E vs. RPPDS-1-E, RPPDS-1-E vs. RPPDS-SCR-E, RPPDS-SCR-E vs. RPPDS-RP-E, RPPDS-0-M vs. RPPDS-1-M, RPPDS-1-M vs. RPPDS-SCR-M, RPPDS-SCR-M vs. RPPDS-RP-M, PPDS-4-E vs. RPPDS-1-E, PPDS-4-M vs. RPPDS-1-M, PPDS-4-E vs. RPPDS-SCR-E and PPDS-4-M vs. RPPDS-SCR-M) to determine whose metabolites contribute significantly to the separation of two samples. Those metabolites with VIP (variable importance in the projection) > 1 and significant changes (*P* < 0.05) in relative contents were differentially expressed between two samples. The metabolite data were also subjected to a combined normalization treatment of log transformation and auto scaling (or pareto scaling alone) before performing PLS-DA.

The metabolites (VIP > 1 and *P* < 0.05) data between the embryos and megagametophytes of PPDS (or RPPDS) were used to produce volcano plot. The volcano plot is a combination of fold change and t-tests. The x-axis is log_2_(FC) in volcano plot. Fold change (FC) was calculated as the ratio of the relative contents of metabolites in embryo to megagametophyte and log_2_ transformed. FC threshold was set as 1.0. Y-axis is -log10 (p.value), and was based on raw p values. Log_2_(FC) value greater than zero indicates that the relative contents of metabolites are higher in embryo than megagametophyte. If log_2_(FC) value is less than zero, this suggest that the relative contents of metabolites are lower in embryo compared with megagametophyte. Metabolic pathway analysis was also performed with metabolites (VIP > 1 and *P* < 0.05) between any of 22 pairs of samples using the MetaboAnalyst web tool to determine whose metabolic pathways are significantly changed. The relative parameters were set using the same method to that used by Song and Zhu (2019).

## ACKNOWLEDGMENTS

We thank Xinghuan Li and Chunxiang Chen for their field support. This study was supported by the National Natural Science Foundation of China (31901300, 31960636) and Natural Science Foundation of Guizhou Province (2019) No. 1165.

## LITERATURE CITED

Ali-Rachedi S, Bouinot D, Wagner MH, Bonnet M, Sotta B, Grappin P, Jullien M (2004) Changes in endogenous abscisic acid levels during dormancy release and maintenance of mature seeds: studies with the Cape Verde Islands ecotype, the dormant model of Arabidopsis thaliana. Planta 219: 479–488

Allen E, Moing A, Ebbels T, Maucourt M, Tomos D, Rolin D, Hooks M (2010) Correlation Network Analysis reveals a sequential reorganization of metabolic and transcriptional states during germination and genemetabolite relationships in developing seedlings of Arabidopsis. BMC Syst Biol 4: 62

Angelovici R, Fait A, Fernie AR, Galili G (2011) A seed high-lysine trait is negatively associated with the TCA cycle and slows down Arabidopsis seed germination. New Phytol 189: 148–159

Arc E, Chibani K, Grappin P, Jullien M, Godin B, Cueff G, Valot B, Balliau T, Jon D, Rajjou L (2012) Cold stratification and exogenous nitrates entail similar functional proteome adjustments during Arabidopsis seed dormancy release. J Proteome Res 11: 5418–5432

Bailly C (2004) Active oxygen species and antioxidants in seed biology. Seed Sci Res 14: 93–107

Baskin JM, Baskin CC (2004) A classification system for seed dormancy. Seed Sci Res 14: 1–16

Bewley JD (1997) Seed germination and dormancy. Plant Cell 9: 1055–1066

Bewley JD, Bradford K, Hilhorst H (2012) Seeds: physiology of development, germination and dormancy. Springer, New York

Bewley JD, Bradford KJ, Hilhorst HWM, Nonogaki H (2013) Seeds: Physiology of development, germination and dormancy (3rd edition). Springer, New York

Bove J, Lucus P, Godin B, Ogé L, Jullien M, Grappin P (2005) Gene expression analysis by cDNA-AFLP highlights a set of new signalling networks and translational control during seed dormancy breaking in Nicotiana plumbaginifolia. Plant Mol Biol 57: 593–612

Cadman CSC, Toorop PE, Hilhorst HWM, Finch-Savage WE (2006) Gene expression profiles of Arabidopsis Cvi seeds during dormancy cycling indicate a common underlying dormancy control mechanism. Plant J 47: 805–822

Carrera E, Holman T, Medhurst A, Dietrich D, Footitt S, Theodoulou FL, Holdsworth MJ (2008) Seed after-ripening is a discrete developmental pathway associated with specific gene networks in Arabidopsis. Plant J 53: 214–224

Chang E, Deng N, Zhang J, Liu JF, Chen LZ, Zhao XL, Abbas M, Jiang ZP, Shi SQ (2018) Proteome-level analysis of metabolism-and stress-related proteins during seed dormancy and germination in Gnetum parvifolium. J Agr Food Chem 66: 3019–3029

Chibani K, Ali-Rachedi S, Job C, Job D, Jullien M, Grappin P (2006) Proteomic analysis of seed dormancy in Arabidopsis. Plant Physiol 142: 1493–1510

Das A, Kim DW, Khadka P, Rakwal R, Rohila JS (2017) Unraveling key metabolomic alterations in wheat embryos derived from freshly harvested and water-imbibed seeds of two wheat cultivars with contrasting dormancy status. Front Plant Sci 8: 1203

De Bont L, Naim E, Arbelet-Bonnin D, Xia Q, Palm E, Meimoun P, Mancuso S, El-Maarouf-Bouteau H, Bouteau F (2019) Activation of plasma membrane H+-ATPases participates in dormancy alleviation in sunflower seeds. Plant Sci 280: 408–415

De Gara L, Paciolla C, De Tullio MC, Motto M, Arrigoni O (2000) Ascorbate-dependent hydrogen peroxide detoxification and ascorbate regeneration during germination of a highly productive maize hybrid: evidence of an improved detoxification mechanism against reactive oxygen species. Physiol Plantarum 109: 7–13

Domergue JB, Abadie C, Limami A, Way D, Tcherkez G (2019) Seed quality and carbon primary metabolism. Plant Cell Environ 42: 2776–2788

El-Maarouf-Bouteau H, Bailly C (2008) Oxidative signaling in seed germination and dormancy. Plant Signal Behav 3: 175–82

Fait A, Angelovici R, Less H, Ohad I, Urbanczyk-Wochniak E, Fernie AR, Galili G (2006) Arabidopsis seed development and germination is associated with temporally distinct metabolic switches. Plant Physiol 142: 839–854

Ferrari F, Fumagalli M, Profumo A, Viglio S, Sala A, Dolcini L, Temporini C, Nicolis S, Merli D, Corana F, Casado B, Iadarola P (2009) Deciphering the proteomic profile of rice (Oryza sativa) bran: a pilot study. Electrophoresis 30: 4083–4094

Finch-Savage WE, Leubner-Metzger G (2006) Seed dormancy and the control of germination. New Phytol 171: 501–523

Galland M, He D, Lounifi I, Arc E, Clément G, Balzergue S, Huguet S, Cueff G, Godin B, Collet B, Granier F, Morin H, Tran J, Valot B, Rajjou L (2017) An integrated “multi-omics” comparison of embryo and endosperm tissue-specific features and their impact on rice seed quality. Front Plant Sci 8: 1984

Gao F, Jordan MC, Ayele BT (2012) Transcriptional programs regulating seed dormancy and its release by after-ripening in common wheat (Triticum aestivum L.). Plant Biotechnol J 10: 465– 476

Joosen RV, Arends D, Li Y, Willems LA, Keurentjes JJ, Ligterink W, Jansen RC, Hilhorst HHW (2013) Identifying genotype-by-environment interactions in the metabolism of germinating Arabidopsis seeds using generalized genetical genomics. Plant Physiol 162: 553– 566

He DL, Yang PF (2013) Proteomics of rice seed germination. Front Plant Sci 4: 1–9

He D, Han C, Yang P (2011a) Gene expression profile changes in germinating rice. J Integr Plant Biol 53: 835–844

He D, Han C, Yao J, Shen S, Yang P (2011b) Constructing the metabolic and regulatory pathways in germinating rice seeds through proteomic approach. Proteomics 11: 2693–2713

Holdsworth MJ, Bentsink L, Soppe WJ (2008a) Molecular networks regulating Arabidopsis seed maturation, after-ripening, dormancy and germination. New Phytol 179: 33–54

Holdsworth MJ, Finch-Savage WE, Grappin P, Job D (2008b) Postgenomics dissection of seed dormancy and germination. Trends Plant Sci 13: 7–13

Hou L, Wang M, Wang H, Zhang WH, Mao P (2019) Physiological and proteomic analyses for seed dormancy and release in the perennial grass of Leymus chinensis. Environ Exp Bot 162: 95–102

Lee CS, Chien CT, Lin CH, Chiu YY, Yang YS (2006) Protein changes between dormant and dormancy-broken seeds of Prunus campanulata Maxim. Proteomics 26: 4147–4154

Liemburg-Apers DC, Willems PHGM, Koopman WJH, Grefte S (2015) Interactions between mitochondrial reactive oxygen species and cellular glucose metabolism. Arch Toxicol 89: 1209–1226

Lin L, Tian S, Kaeppler S, Liu Z, An YQ (2014) Conserved transcriptional regulatory programs underlying rice and barley germination. PLoS ONE 9: e87261

Liu X, Zhang H, Zhao Y, Feng Z, Li Q, Yang HQ, Luan S, Li J, He ZH (2013) Auxin controls seed dormancy through stimulation of abscisic acid signaling by inducing ARF-mediated ABI3 activation in Arabidopsis. Proc Natl Acad Sci USA 110: 15485–15490

Liu Y, Han C, Deng X, Liu D, Liu N, Yan Y (2018). Integrated physiology and proteome analysis of embryo and endosperm highlights complex metabolic networks involved in seed germination in wheat (Triticum aestivum L.). J Plant Physiol 229: 63–76

Lu XJ, Zhang XI, Mei M, Liu Gl, Ma BB (2016) Proteomic analysis of Magnolia sieboldii K. Koch seed germination. J Proteomics 133: 76–85

Müller K, Carstens AC, Linkies A, Torres MA, LeubnerMetzger G (2009) The NADPH-oxidase AtrbohB plays a role in Arabidopsis seed after-ripening. New Phytol 184: 885–897

Murthy UMN, Kumar PP, Sun WQ (2003) Mechanisms of seed ageing under different storage conditions for Vigna radiata (L.) Wilczek: lipid peroxidation, sugar hydrolysis, Maillard reactions and their relationship to glass state transition. J Exp Bot 54: 1057–1067

Narsai R, Gouil Q, Secco D, Srivastava A, Karpievitch YV, Liew LC, Lister R, Lewsey MG, Whelan J (2017a) Extensive transcriptomic and epigenomic remodelling occurs during Arabidopsis thaliana germination. Genome Biol 18: 172

Narsai R, Secco D, Schultz MD, Ecker JR, Lister R, Whelan J (2017b) Dynamic and rapid changes in the transcriptome and epigenome during germination and in developing rice (Oryza sativa) coleoptiles under anoxia and re-oxygenation. Plant J 89: 805–824

Nakabayashi K, Okamoto M, Koshiba T, Kamiya Y, Nambara E (2005) Genome-wide profiling of stored mRNA in Arabidopsis thaliana seed germination: epigenetic and genetic regulation of transcription in seed. Plant J 41: 697–709

Oracz K, Bouteau HEM, Farrant JM, Cooper K, Belghazi M, Job C, Job D, Corbineau F, Bailly C (2007) ROS production and protein oxidation as a novel mechanism for seed dormancy alleviation. Plant J 50: 452–465

Oracz K, El-Maarouf Bouteau H, Bogatek R, Corbineau F, Bailly C (2008) Release of sunflower seed dormancy by cyanide: cross-talk with ethylene signaling pathway. J Exp Bot 59: 2241–2251

Oracz K, El-Maarouf-Bouteau H, Kranner I, Bogatek R, Courbineau F, Bailly C (2009) The mechanisms involved in seed dormancy alleviation by hydrogen cyanide unravel the role of reactive oxygen species as key factors of cellular signaling during germination. Plant Physiol 150: 494–505

Parkhey S, Naithani SC, Keshavkant S (2012) ROS production and lipid catabolism in desiccating Shorea robusta seeds during aging. Plant Physiol Biochem 57: 261–267

Pawłowski TA, Staszak AM (2016) Analysis of the embryo proteome of sycamore (Acer pseudoplatanus L.) seeds reveals a distinct class of proteins regulating dormancy release. J Plant Physiol 195: 9–22

Penfield S, Li Y, Gilday AD, Graham S, Graham IA (2006) Arabidopsis ABA INSENSITIVE4 regulates lipid mobilization in the embryo and reveals repression of seed germination by the endosperm. Plant Cell 18: 1887–1899

Preston J, Tatematsu K, Kanno Y, Hobo T, Kimura M, Jikumaru Y, Yano R, Kamiya Y, Nambara E (2009) Temporal expression patterns of hormone metabolism genes during imbibition of Arabidopsis thaliana seeds: a comparative study on dormant and non-dormant accessions. Plant Cell Physiol 50: 1786–800

Qu CP, Chen JY, Cao LN, Teng XJ, Li JB, Yang CJ, Zhang XL, Zhang YH, Liu GJ, Xu ZR (2019) Non-targeted metabolomics reveals patterns of metabolic changes during poplar seed germination. Forests 10: 659–171

Ren XX, Xue JQ, Wang SL, Xue YQ, Zhang P, Jiang HD, Zhang XX (2018) Proteomic analysis of tree peony (Paeonia ostii ‘Feng Dan’) seed germination affected by low temperature. J Plant Physiology 224–225: 56–67

Rosental L, Nonogaki H, Fait A (2014) Activation and regulation of primary metabolism during seed germination. Seed Sci Res 24: 1–15

Sano N, Permana H, Kumada R, Shinozaki Y, Tanabata T, Yamada T, Hirasawa T, Kanekatsu M (2012) Proteomic analysis of embryonic proteins synthesized from long-lived mRNAs during germination of rice seeds. Plant Cell Physiol 53: 687–698

Sekhon RS, Briskine R, Hirsch CN, Myers CL, Springer NM, Buell CR, de Leon N, Kaeppler SM (2013) Maize gene atlas developed by RNA sequencing and comparative evaluation of transcriptomes based on RNA sequencing and microarrays. PLoS one 8: e61005

Song Y, Zhu JJ (2019) The roles of metabolic pathways in maintaining primary dormancy of Pinus koraiensis seeds. BMC Plant Biol 19: 550

Xia Q, Ponnaiah M, Cueff G, Rajjou L, Prodhomme D, Gibon Y, Bailly C, Corbineau F, Meimoun P, El-Maarouf-Bouteau H (2018a) Integrating proteomics and enzymatic profiling to decipher seed metabolism affected by temperature in seed dormancy and germination. Plant Sci 269: 118–125

Xia Q, Saux M, Ponnaiah M, Gilard F, Perreau F, Huguet S, Balzergue S, Langlade N, Bailly C, Meimoun P, Corbineau F, El-Maarouf-Bouteau H (2018b) One way to achieve germination: common molecular mechanism induced by ethylene and after-ripening in sunflower seeds. Int J Mol Sci 19: 2464

Xu PL, Tang GY, Cui WP, Chen GX, Ma CL, Zhu JQ, Li PX, Shan L, Liu ZJ, Wan SB (2020) Transcriptional differences in peanut (Arachis hypogaea L.) seeds at the freshly harvested, after-ripening and newly germinated seed stages: insights into the regulatory networks of seed dormancy release and germination. PLoS one 15: e0219413.

Xu HH, Liu SJ, Song SH, Wang RX, Wang WQ, Song SQ (2016) Proteomics analysis reveals distinct involvement of embryo and endosperm proteins during seed germination in dormant and nondormant rice seeds. Plant Physiol Biochem 103: 219–242

Yang P, Li X, Wang X, Chen H, Chen F, Shen S (2007) Proteomic analysis of rice (Oryza sativa) seeds during germination. Proteomics 7: 3358–3368

Yazdanpanah F, Hanson J, Hilhorst HWM, Bentsink L (2017) Differentially expressed genes during the imbibition of dormant and afterripened seeds-a reverse genetics approach. BMC Plant Biol 17: 151

Ye N, Zhu G, Liu Y, Zhang A, Li Y, Liu R, Shi L, Jia L, Zhang J (2012) Ascorbic acid and reactive oxygen species are involved in the inhibition of seed germination by abscisic acid in rice seeds. J Exp Bot 63: 1809–1822

Yu Y, Guo G, Lv D, Hu Y, Li J, Li X, Yan Y (2014) Transcriptome analysis during seed germination of elite Chinese bread wheat cultivar Jimai 20. BMC Plant Biol 14: 20

Weitbrecht K, Müller K, Leubner-Metzger G (2011) First off the mark: early seed germination. J Exp Bot 62: 3289–3309

Zaynab M, Pan D, Noman A, Fatima M, Abbas S, Umair M, Sharif Y, Chen SP, Chen W (2018) Transcriptome approach to address low seed germination in cyclobalanopsis gilva to save forest ecology. Biochem Syst Ecol 81: 62–69

Zhou XL, Yang H, Yan QX, Ren A, Kong ZW, Tang SX, Han XF, Tan ZL, Salem AZM (2019) Evidence for liver energy metabolism programming in offspring subjected to intrauterine undernutrition during midgestation. Nutr Metab 16: 20

